# Networks of sexually dimorphic neurons that regulate social behaviors in *Drosophila*

**DOI:** 10.1101/2025.10.21.683766

**Authors:** Gerald M. Rubin, Claire Managan, Marisa Dreher, Elizabeth Kim, Scott Miller, Kaitlyn Boone, Alice A. Robie, Adam L. Taylor, Kristin Branson, Catherine E. Schretter, Adriane G. Otopalik

## Abstract

Neural mechanisms underlying sexually dimorphic social behaviors remain enigmatic in most species. In *Drosophila*, sexually dimorphic P1/pC1x neurons have been described as a site of sensory integration that regulates mating and aggressive behaviors. We show that the male P1/pC1x population forms a highly intertwined network with male-specific mAL and aSP-a neurons that is poised to regulate male behavior. The 48 P1/pC1x cell types exhibit heterogeneous synaptic connections with a subset receiving strong input from identified sensory pathways. We also describe circuit motifs by which P1 and sexually dimorphic aIPg neurons co-regulate social behaviors. Genetic driver lines for these cell types were generated and used to discover distinct roles for P1/pC1x cell types in promoting social acoustic signaling and male-male interactions. Our results reveal unexpected diversity in the connectivity and behavioral roles of the P1/pC1x cell types and provide essential genetic tools for interrogating their neurophysiological and behavioral functions.

## Introduction

Across the animal kingdom, behavior often differs across sexes. Some of the most striking examples have been observed in mating and aggression, wherein different sexes may display distinct behavioral repertoires or contextual dependencies (1–3). The circuit architectures and mechanisms underlying behavioral sex differences remain mysterious in most species, however it is generally thought that sex-specific behaviors arise from sexually dimorphic neural circuits (2, 4, 5).

In *Drosophila melanogaster*, doublesex (*dsx*) and fruitless (*fru*) are key determinants of sexually dimorphic neurons and circuits (5–7). Expression of these two transcription factors dictates post-embryonic survival and development of dimorphic neurons (8–12) required for sexual behaviors (13–17). The dimorphic neurons derived from the pC1 lineage have been the focus of many studies in both males and females (7, 17). The male connectome defines 78 pC1-lineage cells per brain hemisphere in males, which can be grouped into 48 cell types (18). In females, most cells in the pC1 lineage fail to survive and only five cells types, each containing one cell per hemisphere, are found in adults (19, 20). A growing body of literature supports the notion that pC1-lineage neurons integrate conspecific sensory cues (6) and mediate contextually appropriate social behaviors (19, 21–29), balancing courtship and aggression in males (30–35) and receptivity and aggression in females (36–43). All pC1-lineage neurons express *dsx*, but in males these neurons can be divided into *fru*-expressing and non-expressing populations to which distinct functions have been attributed (17, 30). We have adopted the nomenclature of Berg et al. (18) to describe the cells from this lineage. Male-specific cells are called P1 cells and have been divided into 44 cell types, and 19 supertypes in which closely related types have been grouped. Four dimorphic cell types, which we suspect are related to the five pC1-lineage cell types in the female, are called pC1x cell types. This nomenclature takes advantage of the complete inventory of morphologies and synaptic connectivity provided by the connectome, rather than depending upon *fru*- and *dsx*-expression, verification of which is currently incomplete. The extent to which individual P1/pC1x cell types govern different behavioral functions in males is unknown. Precedent for specificity is provided by studies in the female, wherein the five pC1 cell types appear to have different roles in regulating mating and aggressive behaviors (37–39, 41–43).

The other sexually dimorphic neuronal population we examine in detail is the aIPg neurons, which has similar cell numbers in both sexes (18, 44–47). In females, a set of dimorphic aIPg neurons promotes aggressive behaviors (39) by regulating visual pathways, causing a fly to attend to nearby conspecifics (3). Connectomic analysis (45–48) has shown that aIPg and pC1 populations are synaptically connected in the female brain (38, 39, 49). The anatomy and function of distinct aIPg cell types in males, however, remain uncharacterized. Thus, it is unclear how the aIPg–pC1 network shapes male social behavior.

In this report, we describe the synaptic connectivity of the 48 P1/pC1x cell types found in the adult male whole-brain connectome (18). The different P1/pC1x cell types present surprisingly heterogeneous synaptic inputs and outputs, implying that they likely serve distinct circuit and behavioral functions. Strikingly, we found that P1/pC1x neurons form an unanticipated core network with male-specific mAL and aSP-a neurons, two populations previously described to participate in the regulation of social behaviors (19, 25, 28, 44, 50, 51). This highly intertwined network is comprised of nearly 90 male-specific P1/pC1x, mAL, and aSP-a cell types, which together account for nearly one third of all male-specific neurons in the central brain, and is therefore poised to regulate male behaviors. To assess the behavioral functions of the different P1/pC1x neurons in males, we generated 32 new split-GAL4 genetic driver lines covering 34 of the 48 P1/pC1x cell types. We optogenetically activated individual and small subsets of P1/pC1x neurons and provide a first account of how these cell types differentially shape male social acoustic signaling and male-male interactions. Complementing previous work in females (3, 38, 39), we also investigate how P1/pC1x and aIPg neurons co-regulate aggressive behaviors in males. Using a new split-GAL4 genetic driver line, we show that sexually dimorphic aIPg cell types can promote aggressive behaviors in males, likely employing the same downstream circuits that regulate visual attention in females (3). In summary, we provide essential genetic tools, circuit descriptions, and initial behavioral characterizations to enable future neurophysiological and behavioral interrogations of P1/pC1x cell types and the networks in which they reside.

## Results

### Males exhibit 44 P1 and 4 pC1x cell types

During development, pC1 neurons originate from the DM4 dorsal hemilineage (20). Electron microscopic cross-sections of the axonal tract that contain this hemilineage revealed 197 cells in the male (**Figure 1A**), and 125 in the female (**Figure 1B**). This difference likely reflects sex-specific programmed cell death, which prevents most pC1 neurons from reaching adulthood in the female brain (19, 20). Clustering these neurons based on their morphology and connectivity revealed 74 cells per brain hemisphere in the male that had no counterparts in the female. These were first grouped into 44 P1 types (18) which were then collapsed into 19 supertypes by combining closely related cell types. Four other sexually dimorphic cells appear to be related to the five pC1 neuron types (pC1a-pC1e) that survive in females (20) defining four pC1x types (18). The morphologies of these cell types are shown in **Figure 1**. We believe that all 78 cells are likely derived from the pC1 lineage. While higher than early estimates (20, 51, 53) is consistent with recent estimates based on single cell RNA expression analysis (10).

**Fig. 1.**
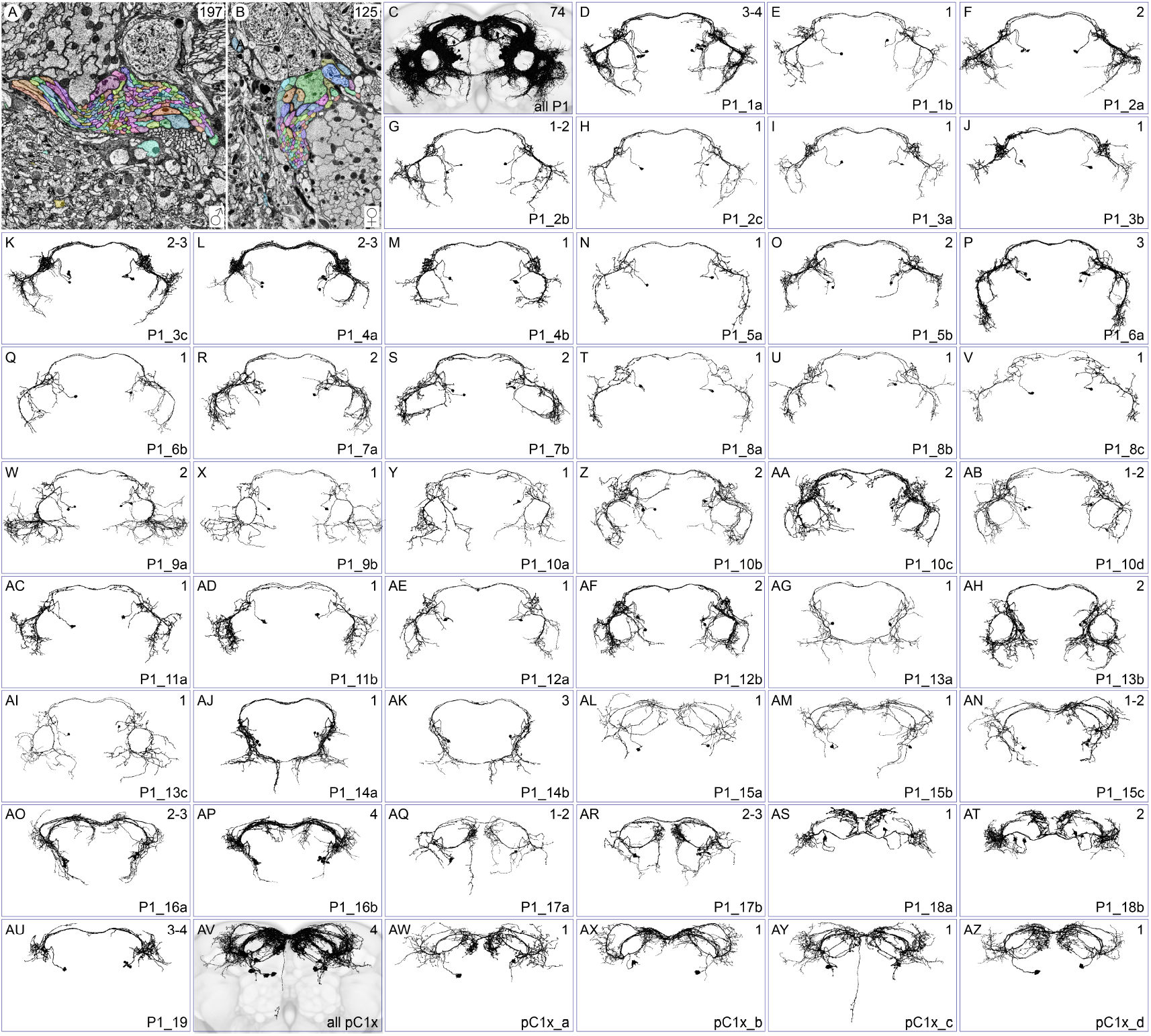
P1/pC1x cell types. (A,B) Cross sections through the dorsal DM4 hemilineage with individual neurons segmented and colored in males and females, respectively. (C) Morphologies of the 74 male-specific P1 cells per hemisphere, superimposed on the JFRC2018 unisex reference brain. (D-AU) Morphologies of the cells comprising individual P1 cell types are shown. The name of the cell type is shown in the lower right and the number of cells of that cell type per hemisphere in the upper right. A range is given when the number of cells observed in the two hemispheres differ; these difference balance out and each hemisphere had 74 P1 neurons. (AV) Morphologies of four sexually dimorphic pC1x cells per hemisphere superimposed on the reference brain (AW-AZ) Morphologies of the individual pC1x cell types. Consistent with previous descriptions using light microscopy (15, 19, 36, 52), all 48 cell types have somata in the posterior medial protocerebrum and have projections to the lateral junction and dorsal commissures.

Important properties of these cell types cannot be resolved solely from connectomic data, such as their expression of *dsx* and *fru*, their neurotransmitter usage, and their neuropeptide expression. Determining these features and assessing the behavioral consequences of activating specific cell types required the development of new genetic tools.

### Generating genetic drivers for P1/pC1x cell types

We generated and characterized split-GAL4 lines that drive expression in individual, or small groups of P1 or pC1x cell types using well-established procedures for intersecting the expression patterns of two enhancers (54). To facilitate these studies, we screened the Janelia FlyLight database (55) for enhancers likely to drive expression in one or more P1/pC1x cell types and used them to generate over 100 new hemidriver lines. Using these new and previously described (54, 56) hemidrivers, we assessed over 1,500 genetic intersections. Those exhibiting specific expression in P1 or pC1x cell types were subsequently established as stable split-GAL4 lines.

To discern which cell types are expressed in each split-GAL4 line, we compared high-resolution confocal image stacks of individual lines to the skeletons derived from electron microscopy (EM) images shown in **Figure 1**. Because we could only use morphological comparisons, these determinations are not as precise as cell-typing in the connectome wherein synaptic connectivity can also be used. As a result, we could not always distinguish closely related subtypes. Moreover, many of our split-GAL4 lines drive expression in multiple cell types. Nevertheless, we were able to produce 32 split-GAL4 genetic driver lines, which allowed us to biochemically characterize and assess the activation phenotypes of roughly half of P1/pC1x cell types. Maximum intensity projection images of these split-GAL4 lines are shown in **Figure 2A-T** and in **Figures S1 and S2**.

**Fig. 2.**
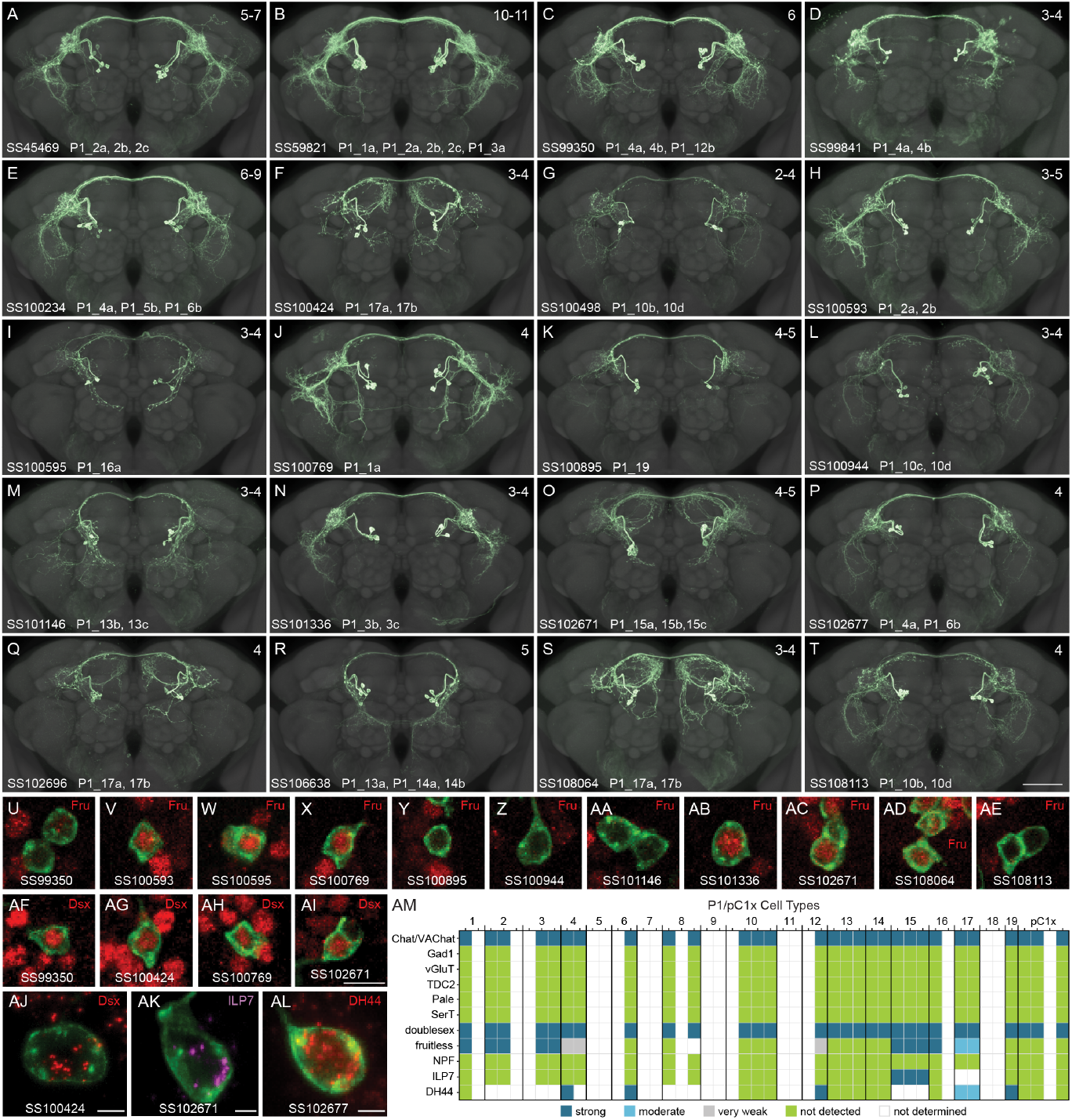
Genetic driver line morphology and expression of key genes. (A-T) Confocal images of the expression patterns in the adult male central brain of individual split-GAL4 lines that express in P1 cell types. Expression patterns are maximum intensity projections shown superimposed on the JFRC2018 unisex reference neuropil. The name of the split-GAL4 line and the cell types in which it drives strong expression are indicated. The number in the upper right of each panel indicates the number of cells per hemisphere with strong expression, based on scoring six brain hemispheres. The scale bar in the lower right of panel T is 50 and applies to panel A-T. The expression pattern of 12 additional split-GAL4 lines used in this work are shown in Figure S1. Figure S2 shows images of expression in each line that include the optic lobes and ventral nerve cord. (U-AI) Confocal images of a small portion of the brain in which the indicated split-GAL4 drives GFP expression (green) and either the male-specific form of the Fru protein (red; U-AE) or Dsx protein (red; AF-AI) has been visualized by antibody staining. (AJ-AL) Images of RNA localization for *dsx*, neuropeptide precursor for ILP7, and neuropeptide precursor for DH44, respectively, obtained using EASI-FISH in 2X expanded brains. Scale bars in panels AI-AL are 5 and the scale bar in AI applies to panels U-AI. Video S1 shows z-stack videos of larger portions of the brains shown in panels U-AL in which multiple cell bodies can be scored. (AM) Matrix summarizing expression of genes encoding neurotransmitter synthesis or transport, Dsx or Fru, or the neuropeptides NPF, ILP7, and DH44 in individual cell types. The light grey indicating very low-level expression of Fru as observed in SS99350 appears to be above background but is far below the level observed in nearby cells in the same preparation and so was scored as negative.

These split-GAL4 genetic driver lines were used in double-label experiments with RNA probes or antibodies against Dsx, Fru, neurotransmitter-related enzymes or transporters, and specific neuropeptides (**Figure 2U-AM and Video S1**). First, we confirmed *dsx* expression, a key defining feature of all P1/pC1x cells, in all 29 of the 48 P1/pC1x cell types that we could directly assess (see examples in **Figure 2 AF-AI** and summary in **Figure 2AM**). We inferred that nine additional cell types were likely Dsx-expressing because other members of their supertype expressed Dsx. For example, we could not assess P1_10a directly because none of our GAL4 lines expressed in that cell type, but found that P1_10b, P1_10c and P1_10d all express Dsx. While we cannot formally exclude the possibility that some of the remaining cell types shown in **Figure 1** lack Dsx expression and thus are not pC1 cells, we use the P1 or pC1x nomenclature for all 48 cell types.

Only a subset of P1/pC1x cell types express *fru*, and it has been proposed that this subset performs distinct behavioral functions (17, 58). Eleven of the 28 cell types we could assess showed Fru expression (**Figure 2U-AE**). Consistent with previous findings (10, 29, 59), all 29 P1/pC1x cell types we could assess utilize the neurotransmitter acetylcholine, and none showed evidence of co-transmission with glutamate, GABA, tyramine, octopamine, serotonin or dopamine. We also assayed expression of three neuropeptides for which prior RNA profiling experiments suggested expression in some P1 cell types (10). We were able to demonstrate the expression of diuretic hormone 44 (DH44) and insulin-like peptide 7 (ILP7), but not neuropeptide F (NPF), in select P1/pC1x cell types (**Figure 2AK-AM**).

### P1/pC1x neurons form a high-level network with mAL and aSP-a neurons

The 48 P1/pC1x cell types project to similar dorsal brain regions but exhibit distinct synaptic inputs and outputs (**Figures S3A, 3A**). The top synaptic connections to the P1/pC1x population at large (constituting > 0.5% of total input or output) are not uniformly connected to all P1/pC1x cell types, yielding the sparse connectivity matrices shown in **Figure 3A**. Notably, most of the strongest connections are with other male-specific neurons (denoted by the suffix ‘m’) situated in the SIP, SMP, and AVLP regions, areas of the brain that exhibit disproportionate innervation by male-specific and sexually dimorphic neurons (18). These include members of the mAL (19, 52) and aSP-a (44) neuronal families (44, 60), two fru-positive populations known to regulate male social behavior (25, 28, 50, 52, 61). Many of the SIP and SMP neuron types among the top P1/pC1x inputs and outputs appear to be members of the aSP-a population (44); 31 cell types have been assigned the aSP-a synonym in neuPrint (18) and we refer to them collectively as the aSP-a cell types.

**Fig. 3.**
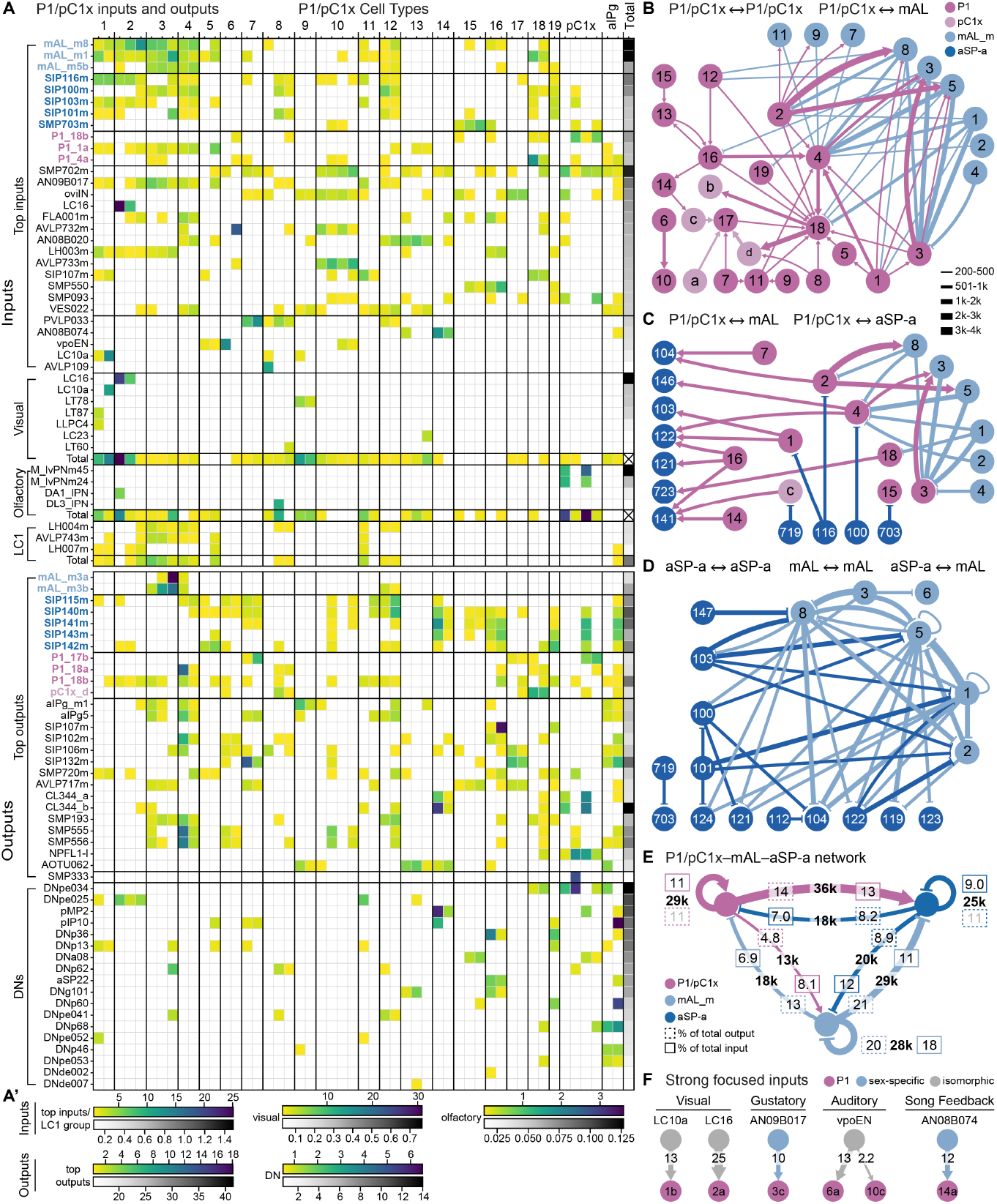
Inputs and outputs of P1 neurons. (A) Top synaptic partners of P1/pC1x cell types. The columns represent the individual P1 and pC1x cell types shown in Figure 1, followed by SMP054-connected and not-SMP054-connected aIPg cell type groups, and finally a column (Total) that represents either the total input provided by that cell type to all P1/pC1x cell types (for Inputs), or total output provided by all P1/pC1x cells to that cell type (for Outputs). Rows represent the indicated cell types whose names are color-coded to match the connectome diagrams in Panels B-E. Rows are presented as sets of input or output types, as indicated, and within a set, related cell types are grouped. Each cell in the matrix shows the strength of a connection represented as a heat map. Scales for the heat maps are shown in panel (A’); heat maps scales differ between sets of rows, as indicated, to allow for the full range of levels to be displayed. Sets of rows are presented in the following order: (1) Top Inputs. The 24 cell types that provide more than 0.5% of the input to P1 cell types taken as a group. For individual cells in the matrix, only connections providing more than 0.5% of that cell type’s input are shown. (2) Five input cell types that, while not meeting the threshold for over 0.5% input to combined P1 types, provide more than 10% of the input to a single cell type. (3) Visual inputs; only connections proving more than 2% of a cell’s input are shown. The last row in this set shows the total amount of visual input to each cell type from all 352 visual projection neurons that connect the optic lobe to the central brain (57). (4) Olfactory input (Olf.). (5) LC1, a group of three cell types that serve as key interneurons connecting pheromone detection and P1 cells. (6) Top Outputs. The 26 cell types that receive more than 20% of their input from P1 cells. SMP333 is shown even though it fails to meet the 20% threshold because of its strong and specific input from pC1x_a. (7) Direct connections to descending neurons (DNs). Figure S3B shows connections to DNs made through a single interneuron and Figure S3C shows the VNC neuropils directly targeted by DNs. (B) P1(purple) and pC1x (light purple) cell types make sparse connections with each other and GABAergic male-specific mAL neurons (light blue). The numbers in the circles represent the P1 or mAL supertypes and a-d represent the pC1x_a – pC1x_d cell types. (C) P1/pC1x cell types make extensive connections with glutamatergic male-specific aSP-a neurons (dark blue). (D) mAL and aSP-a neurons are highly connected. Numbers in circles represent cell types. A key for relating synapse number and line width in B-D is shown. Only connections with >200 synapses are shown in B. Only connections with >500 synapses are shown in C and D; see Figure S4 for diagrams that use a 200-synapse threshold. The full names of the aSP-a cell types are: SIP100m, SIP101m, SIP103m, SIP104m, SIP112m, SIP 116m, SIP119m, SIP121m, SIP122m, SIP123m, SIP124m, SIP141m, SIP146m, SIP147m, SMP703m, SMP719m, SMP723m. (E) Summary diagram of connections between the P1/pC1x, mAL and aSP-a cell type groups. The total number of synapses comprising each connection is shown in the arrow, as well as the percentage of the upstream cell group’s synaptic output dedicated to that connection and the percentage of the downstream cell group’s synaptic input it provides. (F) Examples of strong sensory inputs to individual P1 cell types. Numbers in arrows represent the percentage of the P1 cell type’s input provide by the upstream cell.

Together P1, mAL, and aSP-a form a dense network of male-specific neurons. A subset of P1 neurons is strongly and reciprocally connected with male-specific, GABAergic mAL neurons (**Figure 3B**). Numerous P1 cell types reside up- or downstream of male-specific, glutamatergic aSP-a cell types (**Figures 3C, S4A and S5**). Furthermore, these same mAL and aSP-a neurons form strong connections with each other (**Figures 3D, S4B and S5**).

Connectivity between P1/pC1x cell types is generally sparse. One exception is P1_18 which receives disproportionate synaptic input from other P1 types (**Figure 3B, S5**), raising the possibility that this cell type integrates input from many P1 cell types and conveys P1 population state to downstream neurons, including pC1x_b and pC1x_d. Interestingly, of all the P1/pC1x types, P1_18a specifically provides disproportionate and strong presynaptic input to a male-specific pCd type, SMP710m (see **Figure S4C**). The sexually dimorphic pCd neurons have been reported to regulate the persistence of social arousal and interactions (62).

The overall architecture of this P1/pC1x-mAL-aSP-a network is diagrammed in **Figure 3E** and is more fully described by the connectivity matrix shown in **Figure S5**. While this network diagram illustrates the strong bulk connectivity between these three populations, it masks the sparse connectivity within and across these populations (**Figure S5**); these features of the connectivity will be important for understanding the network’s emergent circuit dynamics. The male CNS contains a total of only 314 male-specific cell types comprised of 1,427 neurons (18); global analyses across the whole brain revealed that these male-specific neurons are preferentially connected with each other and with an additional 924 sexually-dimorphic neurons (18). The P1/pC1x-mAL-aSP-a network is a remarkable example of such preferential connectivity. The 156 P1/pC1x, 98 mAL, and 208 aSP-a neurons in this network are all male-specific and comprise 32% of the male-specific neurons and 28% of the male-specific cell types in the CNS.

### P1/pC1x neuron types receive different sensory inputs

A few P1/pC1x types receive substantial input (more than 10% of total input) from identified sensory pathways (**Figures 3A,F**). The conspecific-detecting visual projection neurons required for courtship and aggressive pursuit, LC10a (3, 63, 64), provide strong input exclusively to P1_1b neurons. The looming-sensitive visual projection neurons implicated in locomotor retreat, LC16 (65, 66), provide remarkably strong presynaptic input to P1_2a neurons, constituting 25% of their total input. The male-specific ascending neurons, AN09B017 (previously referred to as vAB3 (60)), which are responsive to the female-specific pheromone 7,1-heptacosadiene (7,11-HD (25, 28, 67)) via gustatory receptor neurons on the foreleg (25, 28), provide their strongest input to P1_3c neurons and moderate synaptic input to a handful of other P1 cell types. Lastly, the conspecific song-detecting interneurons vpoEN (37) provide 13% of input to P1_6a and moderately strong input to P1_5 and P1_10c. These key sensory pathways provide examples of specific input to distinct P1 cell types.

In contrast, many P1/pC1x types receive second-order chemosensory input from DA1 and DL3, two olfactory receptor neurons (ORNs) responsive to the male-secreted volatile 11-cis-vaccenyl acetate (cVA). When detected by males, cVA suppresses courtship toward female conspecifics (68–70) and promotes aggression toward male conspecifics (71). This social cue is conveyed to several P1/pC1x types via different intermediates. pC1x_a and pC1x_c receive strong input from the second-order antennal lobe projection neurons M_lvPNm24 and M_lvPNm45 (26). Several P1 types receive strong input from LH004m, LH007m, and AVLP743m, collectively referred to as LC1 neurons (30, 72). Other P1/pC1x types receive input from LH003m, comprising part of the population referred to as aSP-f. Lastly, P1_10 neurons receive approximately 5% of their input from neurons that likely correspond to the *fru*-positive tachykinin-expressing cells whose activation can induce aggression (73). Previous work has implicated LC1, aSP-f, and aSP-g as key circuit components in mediating context-dependent, male-specific behavioral responses to cVA (30, 73, 74). Lastly, the mAL neurons also receive strong ascending chemosensory input from the ventral nerve cord (VNC) (25, 28), described in detail by Beckett et al. (companion paper in preparation)(75), and predominantly act upon the P1_1, P1_2, P1_3, P1_4, P1_12, and P1_18 supertypes.

Only a few P1/pC1x neuron types appear to integrate multiple sensory modalities The exceptions are the P1_1, P1_2, and P1_9 supertypes which receive input from both visual and chemosensory pathways.

### P1/pC1x neurons are sparsely connected to descending neurons

Eighteen descending neurons (DNs), the superclass of neurons that connect the brain to the VNC (76– 78), receive strong monosynaptic input from one or more P1, pC1x or aIPg cell types (**Figure 3A**). These downstream DNs include pIP10, DNp13 and pMP2, which are involved in courtship song production (22, 53). P1_16 is upstream of aSP22 (DNa12), which is involved in the coordination of late-stage courtship actions, such as proboscis extension, abdominal bending, and foreleg lifting (79). Altogether, these DNs innervate a diverse set of neuropils in the VNC that control song production and movements of the wing, abdomen, and legs (**Figure S3C**). Seven of the 18 downstream DNs can be matched with *fru*- or *dsx*-positive neurons in the literature (18). The list of unique downstream DNs expands to approximately 30 when disynaptic connections are considered (**Figure S3B**), however these disynaptic connections are likely to have lower effective strength.

### Some P1/pC1x cell types promote courtship song production

We have shown that the different P1/pC1x types in males exhibit heterogeneous synaptic connections. This raises the question of whether they perform distinct circuit and social behavioral functions. During courtship, males extend and vibrate their wings to produce two species-specific songs: pulse and sine. Previous work has implicated descending command-like neurons pMP2, DNp13 and pIP10 in pulse song, sine song, and both pulse and sine song, respectively (22, 53). pMP2 and pIP10 neurons constitute the predominant pathways between the brain and wing neuropil in the VNC. Male-specific P1 neurons, thought to reside upstream of pIP10 and pMP2, are also reported to produce naturalistic patterns of pulse and sine song when activated (22, 80**?**). However, it is unclear which P1 cell types are targeted by the genetic driver lines used in these studies (17, 22, 30, 31, 60).

We leveraged our library of P1/pC1x driver lines, to express the red light-gated cation channel, CsChrimson (81), in single or subsets of P1 or pC1x cell types. We then used a high-throughput recording system equipped with LED boards for optogenetic activation (82) to record evoked song from isolated males in response to 5-second LED stimuli of increasing intensity (**Figure 4A-B**). Using an automated audio classification pipeline (see Methods), we detected bouts of sine and pulse song (**Figure 4B**). Of the P1 cell types activated, none phenocopied pIP10 song production and no pC1x neurons produced song (**Figure 4C**). Only a handful of P1 types produced song during the LED stimulus or post-stimulus periods (**Figure 4C**) and their responses varied in their latency and decay with respect to LED onset and offset (**Figure 4D-H**). For example, co-activation of P1_13a, P1_14a and P1_14b resulted in a complex combination of early pulse song and later sine song epochs during the stimulus period (**Figure 4D**). Activation of P1_17a and P1_17b produced reliable, yet transient, pulse song at LED onset (**Figure 4E**). Activation of P1_19 produced sine song with variable latency and duration during the LED stimulus period (**Figure 4F**). Co-activation of P1_4a, P1_4b and P1_12b, the combination of cell types found in the widely-used P1a split-GAL4 line (31), produced variable patterns of sine and pulse song that persisted well beyond stimulus offset (**Figure 4G**), an activation phenotype consistent with previous observations (80). These diverse evoked responses (**Figure 4H**) suggest that these P1 types may have the capacity to shape song initiation, patterning, and duration.

**Fig. 4.**
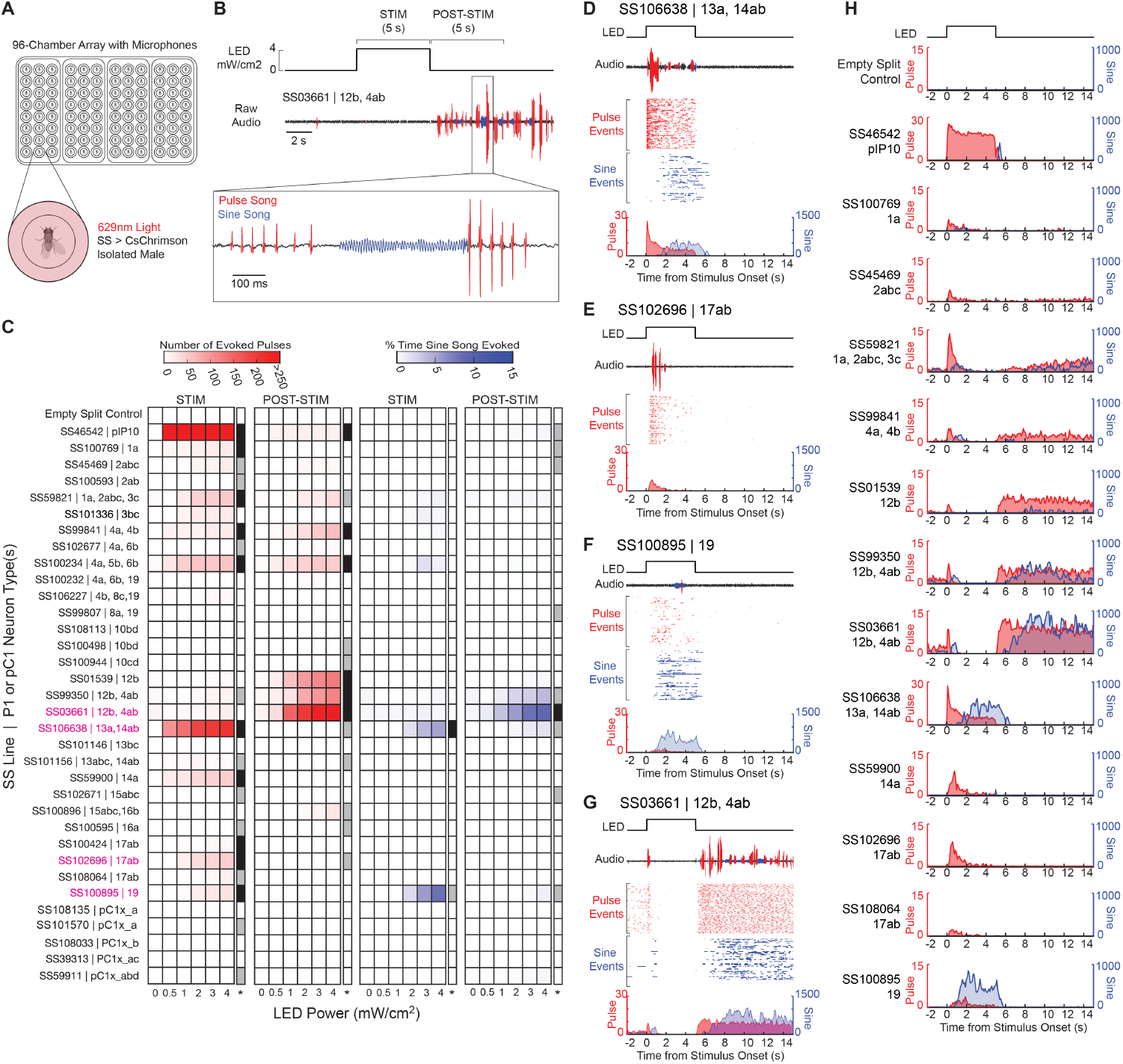
A subset of P1/pC1x cell types promote courtship song. (A) Illustration of high-throughput audio recording system. (B) Representative audio recording of optogenetically-evoked courtship song produced by a single male expressing CsChrimson in P1_12b and P1_4a,b. Bouts of sine (blue) and pulse (red) song were detected during (STIM) and after (POST-STIM) the LED stimulus. (C) Heatmaps showing: mean number of evoked pulses and % time spent producing sine song during STIM and POST-STIM periods across P1/pC1x, Empty (negative control) and pIP10 (positive control) stable split (SS) lines. Means were calculated across four trials at each LED power (N = 16 males for each line). The rightmost column in each heatmap (*), is indicative of the number of LED powers wherein a statistically significant increase in song was observed (gray: significant song observed with highest LED power only; black: multiple LED powers evoked significant song). Statistical significance was determined by one-way ANOVA and Dunnett’s post-hoc test for multiple comparisons (see Methods). (D-G) Song in four representative P1 SS lines evoked by LED (4 mW/cm^2^). For each line: an example audio trace and raster plots show all detected pulse (red) and sine (blue) events across trials (n = 80) are plotted in alignment with the LED stimulus (top). Peristimulus histograms (PSTHs) show the distribution of pulse (red) and sine (blue) event times across all trials with respect to stimulus onset. (H) PSTHs show distributions of pulse and sine event times with respect to stimulus onset (as in D-G) for all lines with notable song phenotypes (indicated by black-shaded cells (*) in C).

Considering the male connectome, P1_5b, P1_7a, P1_7b, P1_14a, and P1_19 are situated directly upstream of pIP10, and P1_14a, P1_14b, and pC1x_c are directly upstream of pMP2, and seven P1 cell types are upstream of DNp13 (**Figure 3A**). Several of the observed song phenotypes could be attributed to these direct connections. For example, P1_19, the only P1 cell type we tested which promotes exclusively sine song, is upstream of pIP10 and DNp13. Prior work showed that co-activation of these DNs results in sine song (53). However, these and several additional P1 types also make disynaptic connections to pIP10, pMP2 and DNp13 (**Figures S3B and Figure 7J**) via different intermediate neurons, complicating the task of mapping these song phenotypes to a wiring diagram. Degeneracy in the circuit is also apparent: activation of different P1 neurons yield similar behavioral outputs. Curiously, similar transient pulse song epochs at LED onset were observed for multiple P1 cell types (e.g. lines targeting P1_4a, 4b, and 12b, P1_14a, or P1_17a and 17b, **Figure 4H**), despite variations in their synaptic connectivity to pIP10 and/or pMP2; as some make monosynaptic connections (e.g. P1_14a, **Figure 3A**) and others make disynaptic connections (e.g. P1_17a, **Figure S3B**). Interestingly, a key song-promoting cell type, P1_14a, also receives strong input from auditory processing pathways and feedback from song-generating motor circuits in the VNC via ascending neuron AN08B074 (**Figure 3F**, see also Figure 5 in Berg et al. (18)), providing a plausible circuit mechanism for prior studies showing that courtship song perception enhances male song production (83–86).

**Fig. 5.**
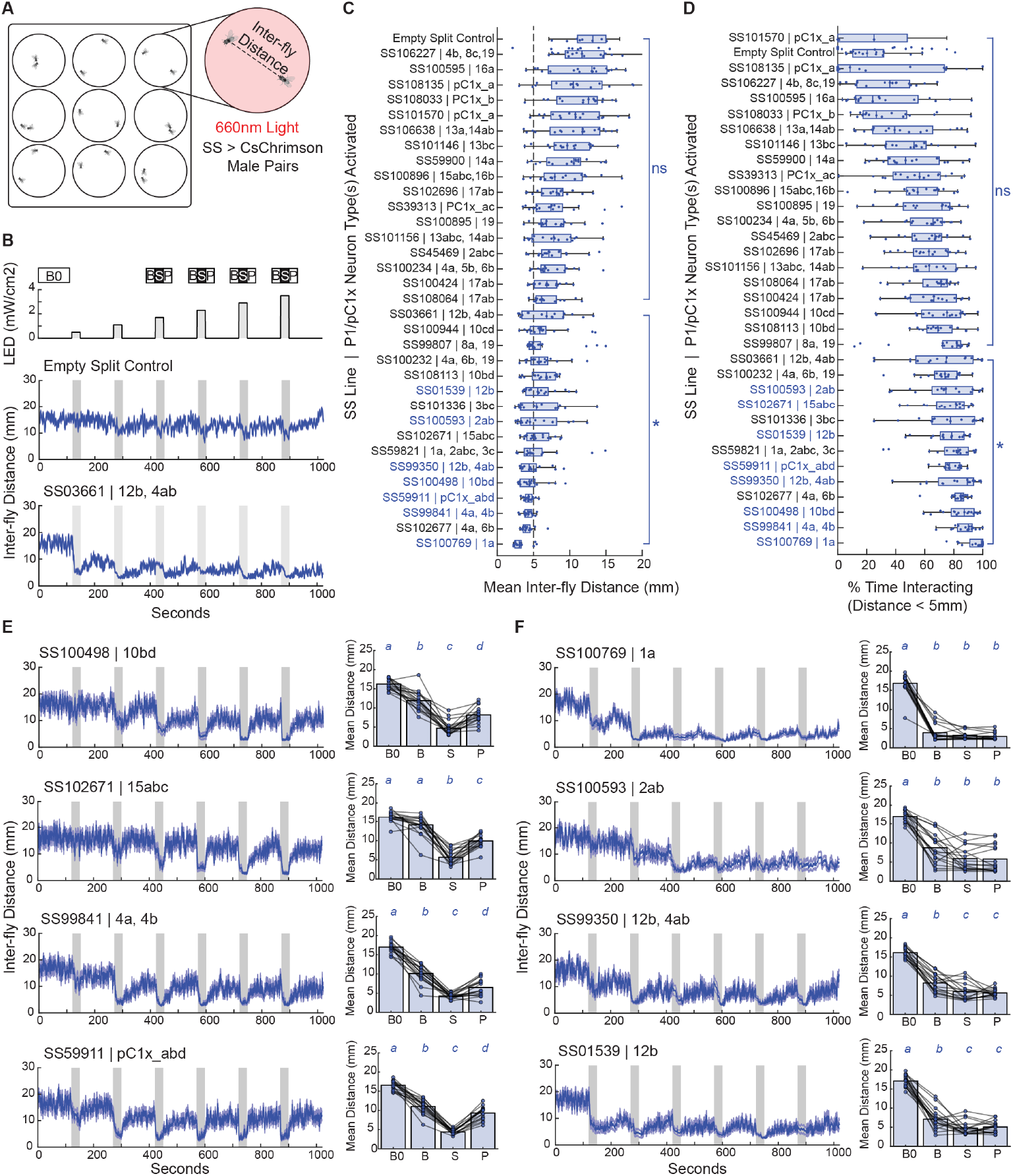
A subset of pC1 neurons promote reliable social interactions. (A) Illustration of nine-chamber platform used for video recording of male-male social interactions. Two male flies of the same genotype (expressing Empty (negative control), P1, or pC1 split-GAL4 lines and UAS-CsChrimson) were loaded into each chamber. (B) Top: All flies were stimulated with 30-second pulses of 660 nm light with two-minute rest periods between stimuli. Middle and bottom: mean inter-fly distances (blue line) with standard error (gray line) over time for male pairs with CsChrimson expression driven by Empty Split-GAL4 (negative control) and a P1 SS line (SS03661) targeting P1_12b and P1_4a,b cells. Gray shaded periods indicate LED stimulus periods. (C-D) Box plots showing the median, upper and lower quartiles of the mean inter-fly distances and mean percent time interacting. For each male-male pair (N = 18 per line), these metrics were calculated across four stimulus periods of highest LED power and are shown as individual data points. The bracket denoted with an asterisk (*) indicates P1/pC1x SS lines that yielded inter-fly distances or % time interacting that are significantly different (p < 0.05; see Methods) from the Empty split-GAL4 negative control animals. (E) Left: mean inter-fly distances (blue line) with standard error (gray line) over time for selected lines showing interactions that are time-locked to LED stimulation periods (shaded in gray). Right: Bar plots showing mean inter-fly distances across male pairs during initial baseline (B0), pre-stimulus baseline (B), stimulus (S), and post-stimulus (P) periods (shown above LED timeseries in B) and for individual pairs (individual data points and line plots). Italicized letters above each bar indicate statistically distinguishable group assignment (p < 0.05). (F) Plots formatted as in E illustrating P1/pC1x lines with social interactions that persist beyond the LED stimulation period (S). This result is supported by assignment of S and P periods to the same statistical groups (see Methods).

Activation of some P1 types resulted in song production at LED offset that persisted well beyond the stimulus period (e.g., the combination of P1_4a, 4b and 12b). Although the circuit mechanisms underlying this sustained song phenotype are difficult to discern from the connectome alone, and the persistent state may be maintained by other neurons (62), it is conceivable that connections between these P1 types and mAL or aSP-a neurons (**Figures 3C and S4A**) play a role in perpetuating the behavior beyond the photoactivation period. Future neurophysiological interrogation during spontaneous and evoked song production will provide deeper insights regarding how the intrinsic and synaptic properties of individual P1/pC1x cell types and P1/pC1x-mAL-aSP-a network dynamics shape natural song initiation and patterning.

### Some P1/pC1x cell types promote male-male social interactions

Acute activation of P1a neurons (P1 types 4a,b and 12b) is sufficient to drive enduring aggressive interactions with other males31 and locomotor pursuit toward moving fly-sized objects (23, 25, 27). It is unknown whether the many other pC1 types in the male promote social interactions.

To test this, we quantified social interactions during acute optogenetic activation of selected pC1 cell types in pairs of males (**Figure 5A-B**) situated in 30mm-diameter chambers. In these conditions, males may interact transiently but maintain an average distance between 10-20mm (**Figure 5B**). Upon optogenetic activation, lower inter-fly distances were observed in approximately half of the pC1 driver lines tested (**Figure 5C**) indicating that some, but not all pC1 cell types, promote social interactions. Thirteen of these 16 lines spent more time interacting at a close distance (<5 mm) than control animals (**Figure 5D**), which is consistent with direct physical contact. Eight of these 16 lines are highlighted in **Figure 5E-F and Video S2**. It is evident in **Video 2** that different P1 types drive different social behaviors. Co-activation of pC1x_a,b,d promotes highly aggressive interactions during the stimulus period, including boxing, lunging, tussling, and wing flicks. This is consistent with prior work using this same line, which implicated these neurons in production of wing flicks in males during competitive courtship (33). In contrast, activation of either P1_1a, P1_2a,b, or P1_15a,b,c appears to drive predominantly chasing and wing extensions, reminiscent of courtship. Lastly, activation of activation of P1_12b or P1_4a,b, or co-activation of P1_12b and P1_4a,b result in wing extensions during the stimulus period, followed aggressive tussling at stimulus offset (consistent with prior observations (31)).

Additionally, the different P1/pC1x cell types promote social interactions that vary in their latency and decay with respect to stimulus onset and offset. Some lines produced tight interactions that are time-locked to the LED stimulation period (**Figure 5E**), and others promoted interactions that persisted well beyond the stimulus period (**Figure 5F**). P1_1a and P1_2ab present the most remarkable persistent phenotypes (**Figure 5F**). Notably, these two P1 types receive direct visual input from LC10a and LC16, respectively (**Figure 3A**), as well as olfactory input from LH003 (aSP-f), which is thought to indicate the presence of the pheromone cVA (74).

Consideration of the two behavioral assessments reveals some themes. The different P1/pC1x types promote varying combinations of song and close interactions. Some P1/pC1x types promote one behavioral class and not the other: P1_13, P1_14, P1_17, and P1_19 promote song, while P1_1a, P1_2a and 2b, P1_10b-d, P1_15a-c, and pC1x_a, b, and d predominantly promote close interactions. P1_4a, 4b, and 12b and a combination of P1_1a, P1_2a-c, and P1_3c promote persistent song and social interactions; these are also the P1 types that connect most strongly to mAL neurons (**Figure 3C**).

### aIPg neurons can induce aggressive behaviors in males

Next, we investigated cell types derived from the fru-expressing aIPg lineage. Our interest in aIPg neurons arose by a chance observation that a split-GAL4 line expressed aIPg1 and aIPg2 in females neurons induced aggression when activated (39). We found that aIPg1 and aIPg2 were the only sexually dimorphic aIPg cell types that provide strong input to SMP054. This isomorphic neuron is a key node in a circuit that promotes social interaction (3). We reasoned that if aIPg neurons were to induce aggression in males by similar mechanisms that they would also connect to SMP054. Using the connectome, we identified three male-specific and four dimorphic aIPg cell types that collectively provide 25% of SMP054’s input (**Figure 6A-H**). There is one male-specific and three dimorphic aIPg neurons in males that are not connected with SMP054 (**Figure 6I-M**), suggesting that these types contribute to other phenotypes.

**Fig. 6.**
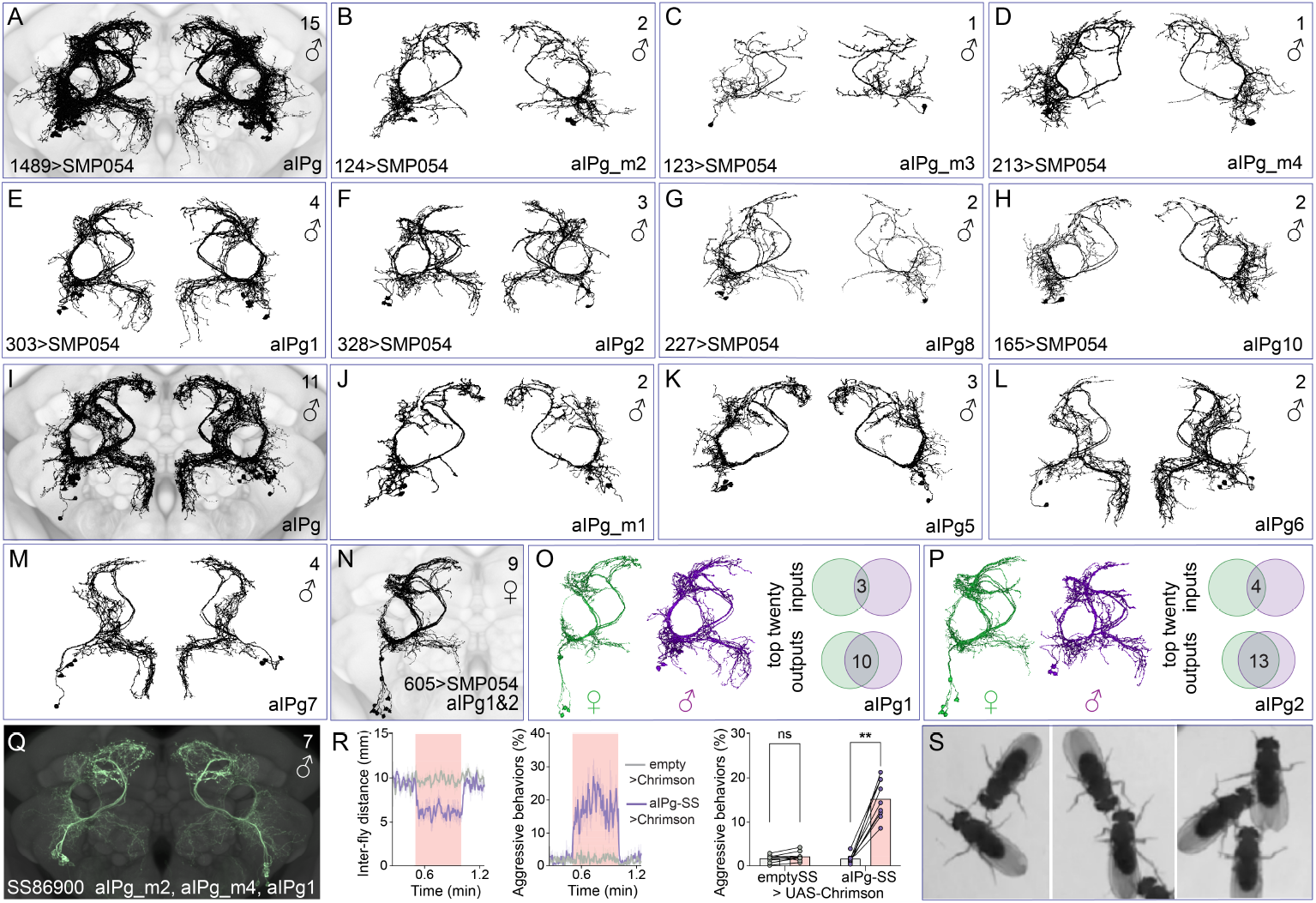
aIPg neurons in males. (A) A composite of the morphologies, derived from EM reconstructions, of three male specific and four sexually dimorphic SMP054-connected aIPg cell types in males shown superimposed on the JFRC2018 unisex neuropil reference. (B-H) Images of the morphology of individual cell types are shown. The number of cells per type in each hemisphere is shown in the upper right of each panel, the cell type name in the lower right, and the number of synapses made to SMP054 in the lower left. In addition to these cells, there is one sexually isomorphic aIPg cell type, aIPg4, that also makes connections to SMP054. (I) A composite of the morphologies of one male-specific and three cell types of sexually dimorphic aIPg neurons that do not connect strongly with SMP054 and (J-M) show their individual morphologies. (N) A composite of the morphologies of the aIPg1 and aIPg2 cell types of sexually dimorphic aIPg neurons in females. These two cell types provide strong input to SMP054 in females and, along with aIPg3, which is female-specific, are the cell types present in the split-GAL4 lines used to study aIPg’s role in female aggressive behaviors (39). (O) Comparison of the sexually dimorphic aIPg1 cell type in males and females. Both the morphologies of these cells differ across sexes as does their connectivity; only three of the top 20 synaptic inputs to aIPg1 are common across sexes, while only ten of their top 20 outputs are in common. (P) Similar comparison of aIPg2 in males and females. (Q) Expression pattern of split-GAL4 line SS86900 in males that express in the three SMP054-connected cell types aIPg_m2, aIPg_m4, and aIPg1; see Figure S2 for an image that includes the optic lobes and ventral nerve cord. (R) Optogenetic activation of SS86900 with 660 nm light at 2.7 mW/cm^2^ results in a decrease in inter-fly distance and an increase in aggressive behaviors. (S) Images of male flies interacting after optogenetic activation of SS86900; images were taken from Video S3.

**Fig. 7.**
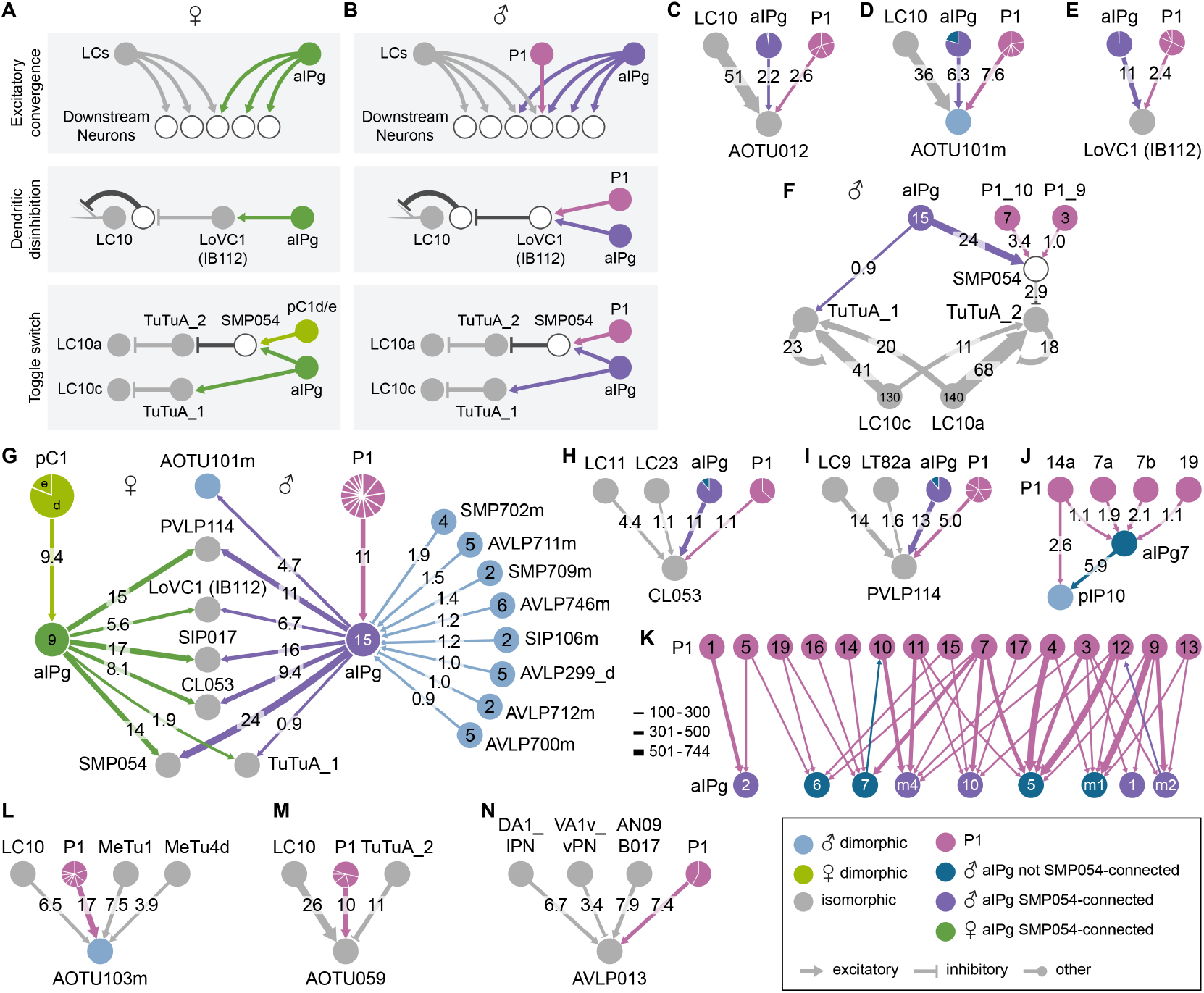
P1 and aIPg neurons coordinately regulate aggression-promoting circuits. (A,B) Circuits regulating the flow of visual information in females (3) and males, respectively. (C,D) LC10, aIPg and P1 neurons all provide input to AOTU012 and AOTU101m and serve as examples of excitatory convergence; the numbers in the arrows indicate the percentage of AOTU012’s or AOTU101m’s synaptic input that comes from each of these cell type families. The sectors in the circles represent individual P1 cell types. Figure S6B shows the positions of these synapses on the arbors of AOTU012 and AOTU101m. (E) In males, P1 neurons participate along with aIPg neurons in the activation of LoVC1 (IB112) which provides dendritic disinhibition of LC10 neurons. (F) Diagram of the toggle switch mechanism in males; compare with Figure 5A of Schretter et al (2025) (3) that shows the same circuit in females. In both sexes, sexually dimorphic aIPg cell types provide strong input to SMP054. However, in males the male-specific P1 types P1_9 and P1_10 take the place of the female-specific pC1d and pC1e. Other aspects of the circuit appear to be isomorphic. (G) Diagram of the inputs and outputs to the SMP054-connected dimorphic aIPg neurons in males and females. In F and G, the numbers in the circles show the number of cells comprising that type and the numbers in the arrows indicate the percentage of the downstream cell’s synaptic input they provide. (H,I) Two of the aIPg downstream cell types that get strong visual input across sexes, CL053 and PVLP114, also get P1 input in males. (J) P1 and aIPg cell types provide input to pIP10. (K) Diagram of the connections between aIPg and P1 neurons with a strength of more than 100 synapses are shown; line widths indicate synapse number. (L) AOTU103m, one of only five male-specific AOTU neurons, gets input from VPNs LC10 as well as MeTu1 and MeTu4d, which provide input from the medulla to the central complex, as well as from multiple P1 cell types. (M) AOTU059, an LC10 target, gets strong input from multiple P1 cell types. While AOTU059 lacks the direct aIPg input often seen in AOTU neurons (see Figure S6), it does get over 10% of its input from the inhibitory neuron TuTuA_2, which is itself inhibited through aIPg and P1 inputs to SMP054 (see panel F). (N) AVLP013 receives three distinct pheromone-sensing inputs as well as input from P1_1a and P1_1b.

Two of the six SMP054-connected aIPg cell types appear to be related to the cell types that promote aggression in females (**Figure 6N**) and provide an instructive example of how dimorphic cell types differ across sexes. Dimorphic cell types arising in a *fru*-positive lineage are thought to result from the differential action of the male and female forms of Fru in the homologous neuronal precursors during development (5). These male versions of the aIPg1 and aIPg2 cell types have morphologies and connectivity that are clearly distinct from their female counterparts (**Figure 6O,P**). Finally, GAL4 expression patterns provide direct evidence for differences in gene expression across sexes; many of the split-GAL4 lines that drive expression in aIPg1 and aIPg2 in females show no expression in males (39).

We generated a split-GAL4 line that expresses in three SMP054-connected aIPg cell types in males (**Figure 6Q**). Optogenetic activation of these aIPg cell types induces low-intensity aggressive behaviors, including fencing (87), (**Figure 6R,S**) in addition to chasing (**Video S3**). This result provides an example of dimorphic cell types that act upon isomorphic partners to promote behaviors that occur in both sexes, a circuit motif we explore more extensively in the next section.

### P1 and aIPg neurons co-regulate isomorphic circuit components

We next investigated the circuits by which the aIPg and P1/pC1x lineages coregulate social behavior in males, as previously shown in females. In both sexes, members of these two lineages can induce aggressive behaviors and are directly connected. Here we show that connections between these two lineages are far more extensive in the male. In females, aIPg neurons promote aggression via three circuit mechanisms (**Figure 7A**) that regulate visual pathways, causing the fly to attend to nearby conspecifics (3). pC1d and pC1e participate in one of the three mechanisms (“toggle switch” in **Figure 7A**). The downstream circuits are largely comprised of isomorphic neurons, including SMP054. We had speculated that conspecific detection and aggressive social interactions may be regulated by the same circuit mechanisms in the male48, but were surprised to find that P1 cell types participate in all three circuit mechanisms, in concert with aIPg cell types (**Figure 7B-F, Figure S6**).

Across sexes, the aggression-promoting aIPg neurons upstream of SMP054 have distinct, dimorphic inputs, and common, isomorphic outputs (**Figure 7G**). More than half of the top 20 inputs in males are from male-specific cell types (including P1 cell types), whereas there are only two female-specific inputs in females (pC1d and pC1e). In contrast, eight of the top 10 downstream cell types are shared across sexes. However, in males many of these isomorphic targets also receive input from P1 cell types (**Figures 7E-F, and 7H,I**).

In males, P1 and aIPg neurons co-regulate downstream circuits by two mechanisms. First, many P1 cell types are upstream of aIPg neurons (**Figure 7J,K**). Nine of the eleven male-specific or dimorphic aIPg cell types receive strong input (>100 synaptic connections) from at least two of the 19 P1 cell type supertypes (**Figure 7K**). Second, as suggested above, P1 and aIPg neurons act upon shared downstream targets, many of which are implicated in social behavior (**Figure 7H-J, L-N)**. This co-regulation is not restricted to visual pathways. For example, aIPg and P1 types reside upstream of pIP10, the DN that drives song production (**Figure 7J**). P1 cell types also regulate both visual and pheromone-responsive sensory streams without the direct involvement of aIPg cell types (**Figure 7L-N**). **Figure S7** shows further examples where p1/pC1x neurons alone, or in combination with aIPg neurons, share downstream targets with either visual (**Figure S7A**) or olfactory (**Figure S7B**) projection neurons. Taken together, these connectomic analyses suggest that pC1 and aIPg cell types play a widespread role in co-regulating sensorimotor pathways implicated in social behaviors.

## Discussion

In this study, we provide a connectome-driven characterization of P1/pC1x neurons, circuits, and behavioral functions in the *Drosophila melanogaster* male. We found that the different P1/pC1x cell types have largely distinct synaptic inputs and outputs. The excitatory, cholinergic P1/pC1x neurons form a network with members of two putatively inhibitory cell type groups: mAL and aSP-a. This highly intertwined network contains one third of all male-specific neurons in the brain and is well-positioned to integrate conspecific sensory information and regulate male-specific social behaviors. We also provide the first detailed description of sexually dimorphic cell types derived from the aIPg lineage in males. These aIPg neurons appear to act both downstream of, and collaboratively with, P1 neurons to regulate social behaviors. Finally, we constructed a set of genetic driver lines for P1/pC1x cell types and used them to demonstrate how these cell types, as foreshadowed by their heterogeneous connectivity, can drive different male behaviors.

### A Sexually Dimorphic Social Network

P1/pC1x neurons are preferentially wired to sexually dimorphic neuron types. Our connectomic analyses revealed that the P1/pC1x population is situated 3-4 layers downstream of sensory receptors implicated in social behavioral decisions. This finding is consistent with the global observation that sex-specific and dimorphic neuron types are concentrated at a mean layer of 4.3 in the male brain connectome (see Figure 3 in Berg et al. (18)). Although it is perhaps unsurprising that most of the major inputs and outputs to the P1/pC1x population are either male-specific or sexually dimorphic neuron types (**Figure 3A**), dimorphic synaptic connections only constitute 6.3% of total connections in males (18). Therefore, P1/pC1x neurons are disproportionately connected to other dimorphic neuron types.

Consideration of the cell types that constitute the top inputs and outputs to the P1/pC1x population revealed that P1/pC1x neurons reside in a core, high-order network with two other male-specific neuronal populations, the *fru*-positive mAL and aSP-a neurons. GABAergic mAL neurons make strong reciprocal connections with select P1 types and distinct sets of glutamatergic aSP-a neurons reside pre- and postsynaptic to different P1/pC1x cell types (**Figures 3A-D, S4 and S5**). Moreover, aSP-a and mAL are extensively and reciprocally connected (**Figure 3D, S4 and S5**). This circuit architecture is more complex than anticipated by previous studies, which predominantly focused on feedforward processing of sensory information by these high-order populations and neurophysiological interrogations in the absence of a connectome roadmap. Understanding how the dynamics of this broader network shape male social behavioral repertoires and decision-making will require connectome-informed, neurophysiological interrogation.

Prior to this study, there had been no characterization of aIPg behavioral functions in males. It was therefore unknown whether aIPg neurons also drive aggressive behaviors in males and, if so, whether they do so coordinately with P1 neurons. Here, we demonstrate that a set of aIPg neurons in males is indeed sufficient to drive low-intensity aggressive interactions (**Figure 6Q-S, Video S3**) and describe how aIPg and P1 neurons co-regulate circuits composed of isomorphic neurons that gate social attention and pursuit in both sexes (3) (**Figure 7**). First, several P1 cell types reside upstream of dimorphic aIPg neurons that regulate these visual pathways. Second, P1 and aIPg neurons act upon common downstream neurons situated in many sensorimotor pathways previously implicated in social behavior. We observed that, while most inputs to aIPg neurons in females are isomorphic, a high proportion of inputs to male aIPg neurons are male-specific cell types, providing an interesting case study of how an otherwise isomorphic circuit is adapted for dimorphic behavioral control in males.

### P1/pC1x Circuits for Multimodal Sensory Processing

The heterogeneous synaptic inputs to P1/pC1x cell types, in combination with the sparse connectivity between them, suggest that sensory information transmitted to individual P1/pC1x types may not propagate across the P1/pC1x population and instead remains segregated in parallel streams to downstream neurons. This said, P1_1b and P1_2a present unique and curious cases of multimodal sensory integration. P1_1b receives approximately 13% of its input from LC10a visual projection neurons (VPNs), which have been implicated in social pursuit (3, 63, 64) and P1_2a receives 25% of its input from the LC16 VPNs, which have been implicated in locomotor avoidance (65, 66). These P1 types also receive chemosensory input from LH003m (aSP-f) and mAL neurons, conduits of the pheromones 7-11 HD and cVA, respectively (7, 25, 30, 74). cVA suppresses courtship (68, 69), whereas 7-11 HD promotes courtship in males (88). Taken together, it is conceivable that P1_1b and P1_2a play key roles in locomotor attraction and avoidance in social contexts.

### Male P1/pC1x Neurons Appear to Perform Distinct Behavioral Functions

We found that male P1/pC1x cell types are sparsely connected to fewer than 20 descending neurons (DNs; **Figure 3A**) that innervate different motor neuropils in the ventral nerve cord (**Figure S3C**). This divergence in synaptic outputs suggests that P1/pC1x neurons perform different social behavioral functions. Indeed, of the P1/pC1x types we assessed, some promoted courtship song (**Figure 4**), others tight interactions (**Figure 5**), some promoted both, and others neither. P1 types with strong connections to mAL neurons yielded behavioral responses that lingered well beyond the photoactivation period. It is unclear whether this reflects mAL’s intrinsic capacity to mediate behavioral persistence, or if such crude optogenetic perturbations elicit artifactual circuit dynamics. Nonetheless, it appears that P1/pC1x types can promote different social behaviors and may have distinct capacities to shape behavioral initiation, patterning, and persistence. Approximately half of the P1 subtypes whose activation phenotypes we studied express *fru*, though we did not see an obvious correlation between expression of *fru* and the nature of the evoked behaviors. Likewise, we found that some P1 cell types expressed either the neuropeptide ILP7 or DH44 but only observed a weak correlation with persistent behavioral phenotypes and DH44 expression in cell types P1_4a, P1_12b and P1_19.

Studies of the five pC1 cell types found in females provide precedent for distinct connectivity and behavioral functions among different pC1 neurons. Depending on mating status, pC1a-c neurons regulate receptivity prior to copulation (37) and egg-laying behaviors post-copulation (42). Recent studies have implicated pC1d specifically in female-female aggression (38, 39) and showed that co-activation of pC1d and pC1e is required to yield long-lasting aggression that persists beyond the activation period (41). Thus, it appears that pC1 neurons in females may have distinct functions that shift in a social history- and context-dependent manner. Further work will be required to understand the functions of each of the 48 P1/pC1x cell types in males, as well as how their neuronal activity and circuit dynamics give rise to flexible social actions and decision-making.

## Concluding Remarks

*Drosophila melanogaster* males and females present striking differences in their sexual and social behavioral repertoires. The availability of brain connectomes in both sexes provides an unprecedented, experimentally tractable venue for study of neural mechanisms underlying this axis of behavioral variation. Only about 5% of neurons differ across sexes, simplifying identification of key circuits for initial interrogation. Although some dimorphic cell types are situated close to the sensory and motor peripheries, most dimorphic or sex-specific cell types reside in the central brain and are disproportionately interconnected (18). Here, as a striking example of this organizational principle, we describe how P1/pC1x neurons are embedded in a dense network with two other male-specific cell type families, mAL and aSP-a. This network contains one-third of all male-specific neurons in the central brain and is poised to play key a role in the high-level control of social behaviors. We also present a detailed connectomic description of sexually dimorphic aIPg neurons, which appear to collaborate with P1/pC1x neurons to execute social behaviors. By generating cell type-specific driver lines, we enable the first assessment of social behavioral functions of identified P1/pC1x cell types and provide the tools needed for future behavioral and neurophysiological study of these neurons and the social decision-making circuits in which they reside.

## Acknowledgments

We thank Kaiyu Wang and Barry Dickson for providing the GAL4 lines SS01539, SS45469, SS59821, SS59900 and SS59911. Kosei Sato and Daisuke Yamamoto provided anti-FruM antibodies. We thank Janelia’s Project Technical Resources led by Gudrun Ihrke for assistance: Christina Christoforou, Kari Close and Yisheng He performed brain dissections and immunohistochemistry. Kari Close performed EASI-FISH experiments. Janelia’s FlyLight Project Team and Project Pipeline Support team led by Geoffrey Meissner, especially Allison Vannan, Jennifer Jeter, and Joanna Hausenfluck performed CNS dissections, staining, and imaging. Janelia’s Invertebrate Shared Resource and Scientific Computing contributed to stock generation and image processing. Hideo Otsuna helped identify matches between EM skeletons and light microscopic images of enhancer-driven expression patterns. Geoffrey Meissner and Rob Svirskas contributed to the split-GAL4 website. Heather Dionne generated hemidriver constructs. We thank Carmen Morrow, Erin Solomon, and Jinyang Liu for their critical roles in constructing and supporting the video recording system. We also thank the Janelia Experimental Technology team, particularly Steve Sawtelle and Mioara (Andrea) Gugiu for their roles in constructing electronics components for both behavior systems. We thank David Stern for letting us use his laboratory’s audio recording system for pilot experiments. We thank the Janelia EM Project Team and the Cambridge Connectomics Group, especially Philipp Schlegel and Isabella Beckett, for their extensive input and advice throughout this work. We thank Josh Lillvis and David Anderson for their insightful comments on the manuscript. We thank Aaron Allen, Megan Goodwin, and Stephen Goodwin for helpful discussion, communicating data prior to publication, and comments on the manuscripts. This work was supported by the Howard Hughes Medical Institute.

This article is subject to HHMI’s Open Access to Publications policy. HHMI lab heads have previously granted a nonexclusive CC BY 4.0 license to the public and a sublicensable license to HHMI in their research articles. Pursuant to those licenses, the author-accepted manuscript of this article can be made freely available under a CC BY 4.0 license immediately upon publication.

## Methods and Materials

### Connectome analyses

Our analyses are based on the male CNS dataset (v0.9 (18, 57)) and the female hemibrain dataset (v1.2.1 (45)) as queried using the neuPrint interface (neuprint.janelia.org) unless otherwise noted. A complete list of synaptic connections used to construct our circuit diagrams can be found in neuPrint. Synaptic connections organized by neuron pair and neuropil were obtained from neuPrint (using neuprint-python fetch_adjacencies with default settings). Connection strengths are based on the number of synapses. The connection tables obtained in this way do not include synapses with small, untyped EM bodies and such synapses were not included when calculating relative connection strengths. Including these additional synapses in calculations of relative connection strength (as done in the neuPrint web interface) results in lower numbers that most likely underestimate the relative contribution of one cell type to the inputs or outputs of another type. To quantify the contributions from P1/pC1x neurons to DNs made through a single interneuron, we computed the connection strengths of all such paths. For each path, we multiplied the input-normalized connection strengths (that is, input connection summing up to 1) across the two synapses. Then we summed over all such paths between each pair of P1 or pC1x neuron and the DN to arrive at the heatmap shown in Figure S3B. This procedure is equivalent to raising the input-normalized connectivity matrix to the second power. See Hoeller et al. 2025 (89) for more details. A comprehensive list of P1 cell inputs and outputs is provided in Supplemental File 1.

### Generation of split-GAL4 lines

Split-GAL4 lines were generated as previously described (57, 65, 90–93). Databases of expression patterns generated in the adult brain by single genomic fragments cloned upstream of GAL4 (56, 94) were manually or computationally (55) screened. Individual enhancers that showed expression in the desired cell type were then cloned into vectors that contained either the DNA-binding or activation domain of GAL4 (95, 96). Approximately 100 new constructs of this type were generated specifically to support the current work. These constructs were combined in the same individual and screened for expression in the desired cell type by confocal imaging. Over 1,500 such screening crosses were performed to generate the new split-GAL4 lines reported here. Successful constructs were made into stable lines.

### Characterization of split-GAL4 lines

Stable split-GAL4 lines were characterized by confocal imaging of the entire expression pattern in the brain and VNC at 20X. Most lines were also imaged at higher magnification (63x) and/or subjected to stochastic labelling (MCFO99) to reveal the morphology of individual cells. Split-GAL4 images are shown as MIPs after alignment to JRC2018100. The original confocal stacks from which these images were generated, and other key imaging data can be downloaded from www.janelia.org/split-GAL4. Determining the correspondence between the cell types present in each split-GAL4 line and those described in the connectome (18) was based solely on morphology. Morphologies were compared using the VVDViewer (https://github.com/JaneliaSciComp/VVDViewer/wiki) image comparison tool by comparing high-resolution (63X) confocal stacks that had been aligned to the JFRC2018 unisex template with skeletons of individual P1/pC1x neurons that had been transferred into the same coordinate system. Skeletons for all neurons in the male CNS volume can be downloaded from https://console.cloud.google.com/storage/browser/flyem-male-cns/v0.9/segmentation/skeletons-unisex-template/skeletons-swc. Even when assigning correspondence between cells in two different connectomes, where information on connectivity can also be employed, the process is not always straightforward (47). Due to the morphological similarity among many of the P1/pC1x cell types, assigning correspondence to individual cell types defined by connectome analysis was often challenging and there is a small chance of misassignment between closely related cell types. Some lines exhibited off-target expression in either non-P1/pC1x cell types or displayed very low-level expression in other P1/pC1x cells. Such expression can be seen in the images shown in Figure 2 and Supplementary Figures 1 and 2 or in the original confocal stacks from which these images were derived which can be found at janelia.org/split-GAL4. A further technical limitation is that expression in cell types too faint to be detected in our images, might still contribute to observed activation phenotypes. This issue is mitigated to some extent by using multiple, genetically distinct split-GAL4 lines that express in the same or overlapping cell types as well as by comparing the effects of different levels of activation.

### Immunohistochemistry

The dissections, immunohisto-chemistry, and imaging of fly central nervous systems were done principally as previously described (90) (https://www.janelia.org/project-team/flylight/protocols). In brief, brains were dissected in Schneider’s insect medium and fixed in 2% paraformaldehyde (diluted in the same medium) at 25°C for 55 min. Tissues were washed in PBT (0.5% Triton X-100 in phosphate-buffered saline) and blocked using 5% normal goat serum before incubation with antibodies.

Brains expressing myristolated GFP (JFRC29; as previously described (96)) driven by the indicated split-GAL4 line were stained with chicken anti-GFP (Invitrogen A10262, 1:1000) and either rat anti-FruM (1C1, hybridoma supernatant 1:10; gift from K Sato and D. Yamamoto (97)) or mouse anti-DsxDBD (Developmental Studies Hybridoma Bank, Univ. Iowa, hybridoma supernatant 1:2). Secondary antibodies were Alexa Fluor®Plus 488-conjugated goat anti-chicken (Invitrogen A3293, 1:800) in combination with Alexa Fluor®594-conjugated goat anti-rat (Jackson ImmunoResearch 112-585-167, 1:800) and Alexa Fluor®568-conjugated goat anti-mouse (Invitrogen A11031, 1:400), respectively. After staining and post-fixation in 4% paraformaldehyde, brains were mounted on poly-L-lysine-coated cover slips with the posterior side up, cleared, and embedded in DPX as described. Image z-stacks were collected using an LSM880 upright confocal microscope (Zeiss, Germany) fitted with a Plan-Apochromat 40X/1.3 M27 oil immersion objective. The voxel size was 0.44 x 0.44 x 0.44 mm3 (688 x 688 pixels in xy).

### RNA in situ hybridization

Adult females (5 to 7 days post eclosion) were expanded, probed, and imaged using the EASI-FISH method as previously detailed (93, 98). The oligo probes and HCR hairpins were designed by, and purchased from, Molecular Instruments, Inc. Imaging was performed on a Zeiss Z7 microscope equipped with a 20X objective. Laser power and exposure time were optimized to maximize the signal-to-noise ratio.

### Fly Husbandry for Behavioral Experiments

For optogenetic activation of individual or subsets of P1/pC1x neurons, P1/pC1x split-GAL4, pIP10 split-GAL4 (positive control), and empty split-GAL4 (negative control) male flies were crossed to 20XUAS-CsChrimson-mVenus [attp18] female flies. All crosses and selected progeny were housed in the dark at 18°C and 50% humidity. Crosses were maintained on food supplemented with 0.2mM trans-retinal (Sigma-Aldrich). Male progeny were collected 0-2 days after eclosion and group-housed on standard food supplemented with 0.4mM trans-retinal for 5-7 days prior to behavioral experiments.

### Evoked Song Recordings with Optogenetic Activation

Audio recordings were acquired at 5000 Hz using a replica of a previously published high-throughput recording system (82). In brief: the system is equipped with an array of 96, 5 mm-diameter chambers (Figure 4A). Each chamber is equipped with an isolated microphone at the base and clear acrylic on top. Individual male progeny were transferred from food vials to chambers by aspirator. For optogenetic activation, the chamber array was illuminated with diffuse 625 nm light from above for 5-second periods at increasing intensities of: 0, 0.5, 1, 2, 3, and 4 mW/cm^2^, with four trials at each intensity. A sample size of 16 male flies were tested for each driver line. Recordings were acquired at 25°C and 40% humidity between zeitgeber times (ZT) 3-8.

### Audio Processing and Analysis of Song Recordings

Evoked courtship song was detected and classified in the raw audio recordings for individual males using SongExplorer (https://github.com/JaneliaSciComp/SongExplorer). In brief: a convolutional neural network model was trained to detect pulse and sine song, as well as ambient sounds and noise artifacts (“other sounds”). Ground-truth data for the model consisted of the following manually annotated *Drosophila melanogaster* song events: 613 pulses, 402 sine waves, 294 inter-pulse intervals, 101 ambient waves, and 217 other sounds. The model was trained with four million steps, using a learning rate of 0.000001 until the loss, accuracy, and error rates plateaued (as in Lillvis et al., 2024 (53)). This resulted in precision/recall statistics of 93%95% for pulse song, 96%96% for sine song, 93%94% for inter-pulse intervals, 94%95% for ambient sound, and 95%93% for other sounds. SongExplorer produced event start and end times for each sound class.

The sound classifications were utilized in tandem with the raw audio recordings for plotting and analyses using custom MATLAB scripts. To determine if optogenetic activation of each P1/pC1x driver line promoted song production, two metrics were quantified at each LED intensity for each fly: mean number of evoked pulses and mean percent time producing sine song, during LED stimulation (4mW/cm^2^ 629 nm light from above) and post-stimulation periods (as illustrated in Figure 4B). Across LED intensities, these metrics were compared to a no-stimulation period (0 mW/cm^2^) using a one-way ANOVA and Dunnett’s post-hoc test for multiple comparisons. For selected P1/pC1x driver lines, pulse and sine event times were plotted as LED stimulus-aligned raster plots across trials (Figure 4D-G), spanning from two seconds prior to stimulation to 10 seconds after stimulation. This 17-second period was divided into 100-millisecond bins, and pulse and sine events were counted across these 170 bins to generate mean peristimulus histograms for P1/pC1x lines with significant song phenotypes (Figure 4H).

### Male-Male Behavior Experiments with Optogenetic Activation

For male pairwise experiments: Video recordings of male pairs were acquired at 60 frames per second and 1100 × 1100-pixel resolution (at 8.5 pixels per millimeter) on a custom behavioral rig intended for stability and high-throughput behavioral experiments. In brief: a monochrome Blackfly USB camera (FLIR) with a 25mm lens (Edmund Optics 86-572) and a UV/VIS cut filter (Edmund Optics 89-836) was positioned above a 3×3 array of 30 mm-diameter chambers. The chambers are composed of a custom-printed, flat, white base that sufficiently transmits visible and infrared wavelengths and clear, vacuum-formed, plastic circular lids. For optogenetic stimulation and backlighting, a custom LED board (https://github.com/janelia-experimentaltechnology/RGB-IR-LED-Boards/tree/main) equipped with an array of 660 nm, 465 nm, and 860 nm LEDs was situated below the chambers and an acrylic diffusion layer. The on-off status of the LEDs used for optogenetic stimulation was indicated by an 860 nm LED within the camera field of view and monitored to ensure the stimulation protocol was properly executed for every experiment. The system is controlled with custom MATLAB software.

Pairs of males were loaded into chambers by aspiration. During the experiments, the chambers were illuminated with continuous low-level 860 nm light (1 mW/cm^2^) and 465nm light (0.1 mW/cm^2^) for video capture and to ensure the flies were visible to each other, respectively. For optogenetic activation, 660 nm light was emitted for 30-second periods at increasing intensities of: 0, 0.5, 1.1, 1.7, 2.3, 2.9, and 3.5 mW/cm^2^ with 2-minute rest periods between stimuli. Two recording sessions were conducted for each line, resulting in a sample size of 18 male pairs for each driver line. The previously published P1a (SS03661 (31)) and empty split driver lines were used as positive and negative controls, respectively, and recorded once per recording session. Recordings were acquired at 25°C and 40% humidity.

Video was captured at 60 FPS and post-hoc tracking of fly x-y coordinates enabled the calculation of inter-fly distances. Data were analyzed using one-way ANOVA with Bonferroni post-hoc test for multiple comparisons.

### Video Processing and Analysis of Male-Male Behavior

The videos were processed using the Fly Disco Analysis pipeline (99), available at https://github.com/kristinbranson/FlyDiscoAnalysis, with new functionality to handle videos of multiple chambers. In brief: fly coordinates were tracked using Caltech FlyTracker (100) (available at https://github.com/kristinbranson/FlyTracker) and registered on a per-chamber basis. Custom MATLAB scripts were used to calculate two metrics for each male-male pair: mean interfly distance and mean percent time interacting (when the two flies are within 5mm of each other), for four experimental periods: a 120-second period prior to any stimulation (B0), across the four highest-power LED stimulation periods (S; 120 seconds total), across four 30-second post-stimulation periods following four highest-power LED stimuli (P; 120 seconds total), and the four 30-second pre-stimulation periods preceding the four highest-power LED stimuli (B; 120 seconds total). These experimental periods are shown in Figure 5B.

To determine if optogenetic activation of each P1/pC1x driver line promotes social interactions, mean inter-fly distances and mean percent time interacting during the stimulation (S) periods were compared across all driver lines (Figure 5C-D) using a one-way ANOVA and Bonferroni post-hoc test for multiple comparisons. Mean inter-fly distances with standard error margins were plotted over time for selected P1/pC1x driver lines with significant interaction phenotypes (Figure 5E-F). Mean inter-fly distances during each of the four experimental periods (B0, B, S, P) were plotted for all male pairs for each line and comparisons between these experimental periods were assessed using a one-way ANOVA and Bonferroni post-hoc test for multiple comparisons.

### Group Behavioral Experiments with Optogenetic Activation of aIPg

Groups of 8 males were recorded in a 53.3 mm x 3.5 mm circular arena previously detailed (3, 99). In brief: all tests were conducted under visible light conditions using white-light illumination from above (7 /cm^2^) and flies were loaded into the arena using an aspirator. Videos were recorded from above using a camera (USB 3.1 Blackfly S, Monochrome Camera; Point Gray, Richmond, Canada) with an 800 nm long pass filter (B and W filter; Schneider Optics, Hauppauge, NY) at 170 frames per second and 1024 × 1024-pixel resolution. For optogenetic activation, the arena was illuminated with a constant 860 nm LED light (6.2 mW/cm^2^) and 660 nm light was emitted for three 30-second periods at 2.7 mW/cm^2^ with 30-second rest periods between stimuli. Recordings were acquired at 25° C, 60% humidity, and between ZT0 – ZT4.

### Video Processing and Analysis of Group Behavior

Following collection, videos were processed using the Fly Disco Analysis pipeline (99). In brief: flies were tracked using Caltech FlyTracker (100) and accessed at https://github.com/kristinbranson/FlyTracker) followed by automated classification of behavior with JAABA classifiers (101). A novel classifier for male aggression was created based on prior definitions and excluded unilateral wing extension. The performance of these classifiers was validated against manually labeled ground truth data using videos that were not part of the training dataset (Framewise performance: 90.5% true positive, 86.8% true negative). Aggression was computed on a per-arena basis rather than a per-fly basis using a previously detailed MATLAB script (3). The percent aggression is calculated as the sum of the aggression scores for all the trajectories wherein flies exhibited aggression divided by the total number of flies in the arena and multiplied by 100. Behavioral time courses display the mean of 0.35 s (60-frame) bins. For Figure 6R, only the first stimulus period (2.7 mW/cm^2^) is shown out of the six-stimulus protocol. Comparisons between experimental periods were assessed using a Wilcoxon matched-pairs signed rank test.

---

**Fig. S1.**
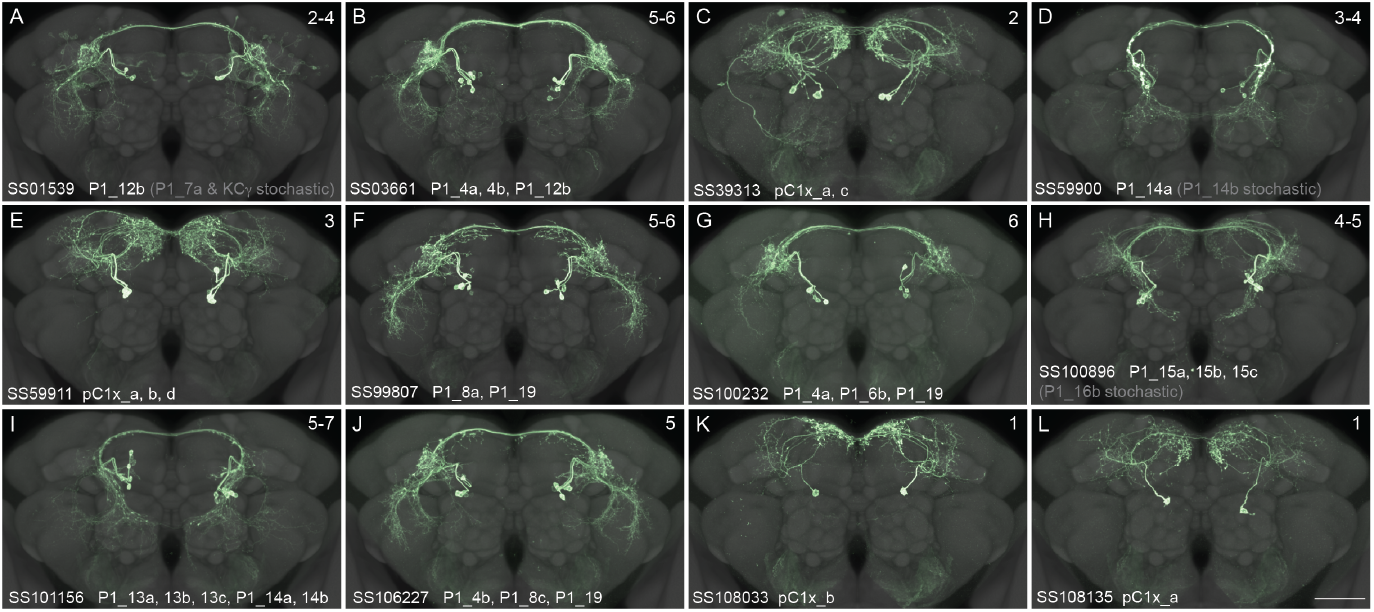
Additional split-GAL4 lines for P1 and pC1x cell types. (A-L) Confocal images of the expression patterns in the adult male central brain of individual split-GAL4 lines that express in p1/pC1x cell types. Expression patterns are maximum intensity projections derived from 63X confocal stacks and are shown superimposed on the JFRC2018 unisex reference neuropil. The name of the split-GAL4 line and the cell types in which it drives strong expression are indicated. The number in the upper right of each panel indicates the number of cells per hemisphere with strong expression, based on scoring six brain hemispheres. The scale bar in the lower right of panel L is 50and applies to panel A-L.

**Fig. S2.**
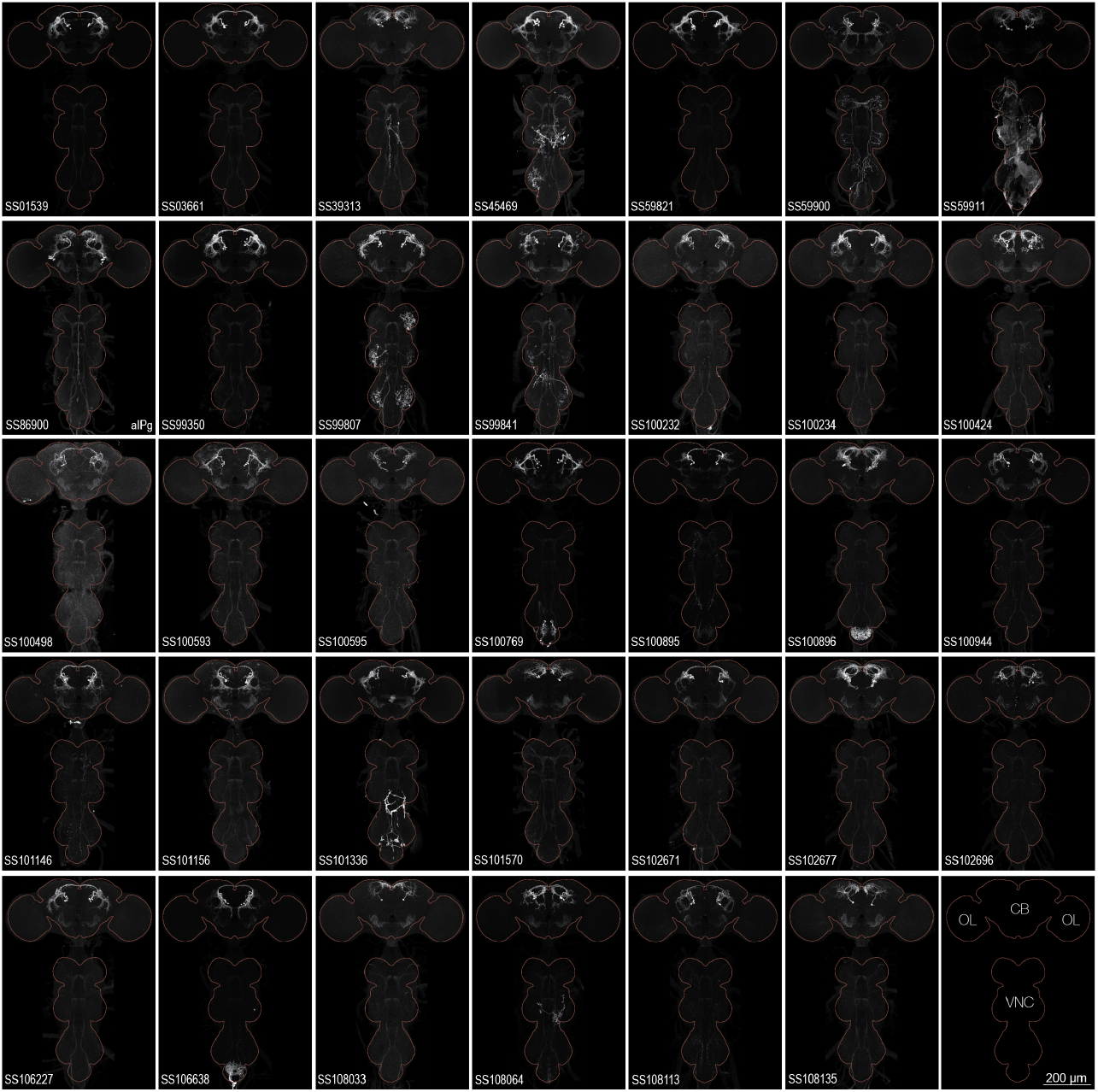
Images of split-GAL4 lines showing central brain, optic lobes, and ventral nerve cord. Maximum intensity projections (20X confocal images) of the expression patterns driven by stable split-GAL4 lines for the indicated 34 lines. Brains and VNCs are outlined in red. The lower right panel shows a key with the central brain (CB), optic lobes (OL) and ventral nerve cord (VNC) indicated, and a scale bar that applies to all panels.

**Fig. S3.**
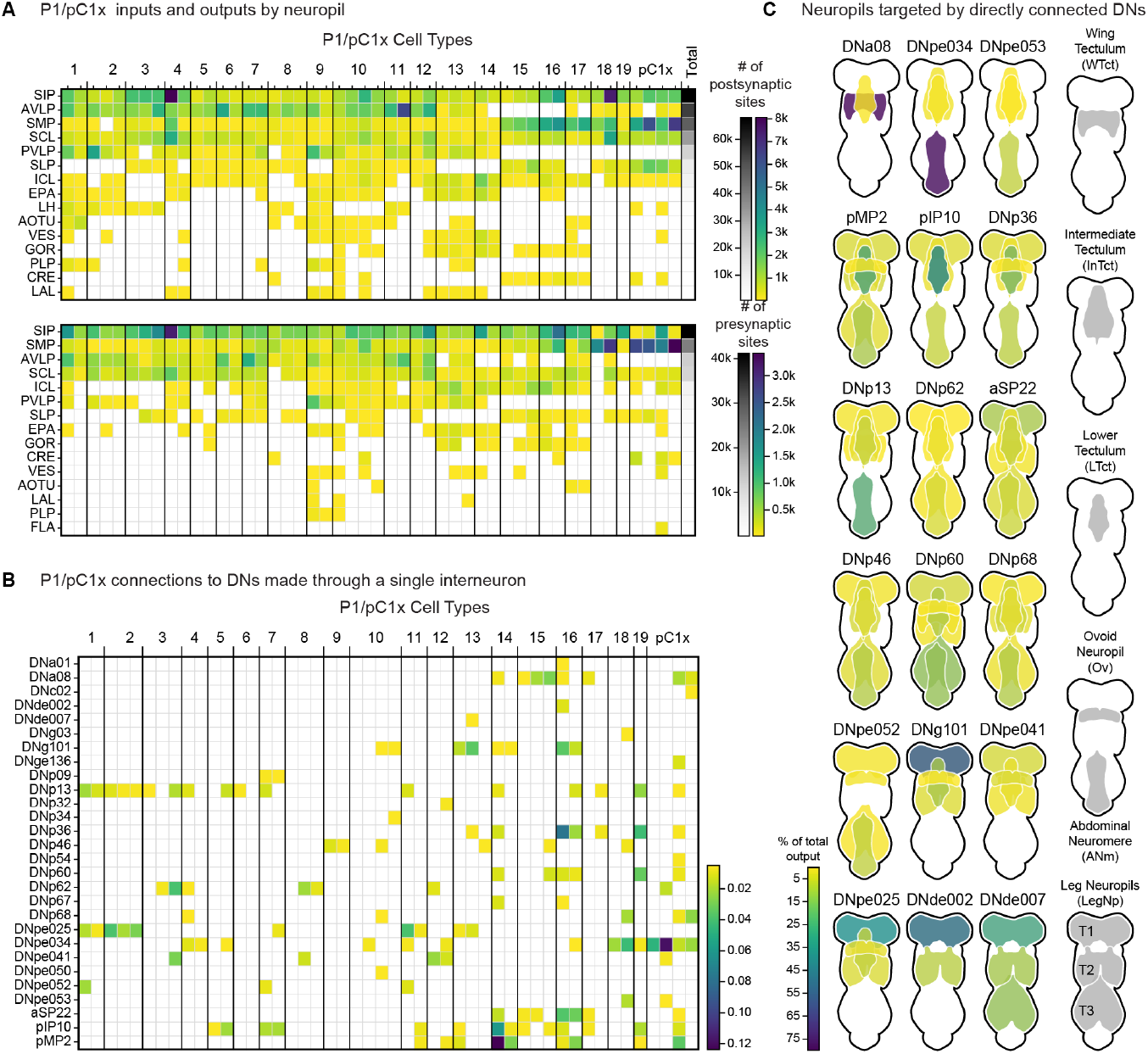
Additional matrices of inputs and outputs of P1 neurons. (A) Brain regions providing input or receiving output from the indicated cell types. The fill of each cell is a heatmap indicating the fraction of that P1 or pC1x cell type whose input (estimated by synapse number) comes from the indicated brain region (Inputs) or the fraction of its output that goes to that brain region (Outputs) (B) Inputs to DN cell types provided by P1 and pC1x neurons via a single interneuron. The heat-map scale represents propagated connection strengths (see Methods). (C) Schematic of VNC neuropils targeted by each of the DNs that are directly downstream of P1s (see Figure 3A). Heat map represents the percentage of that DN’s output going to each neuropil.

**Fig. S4.**
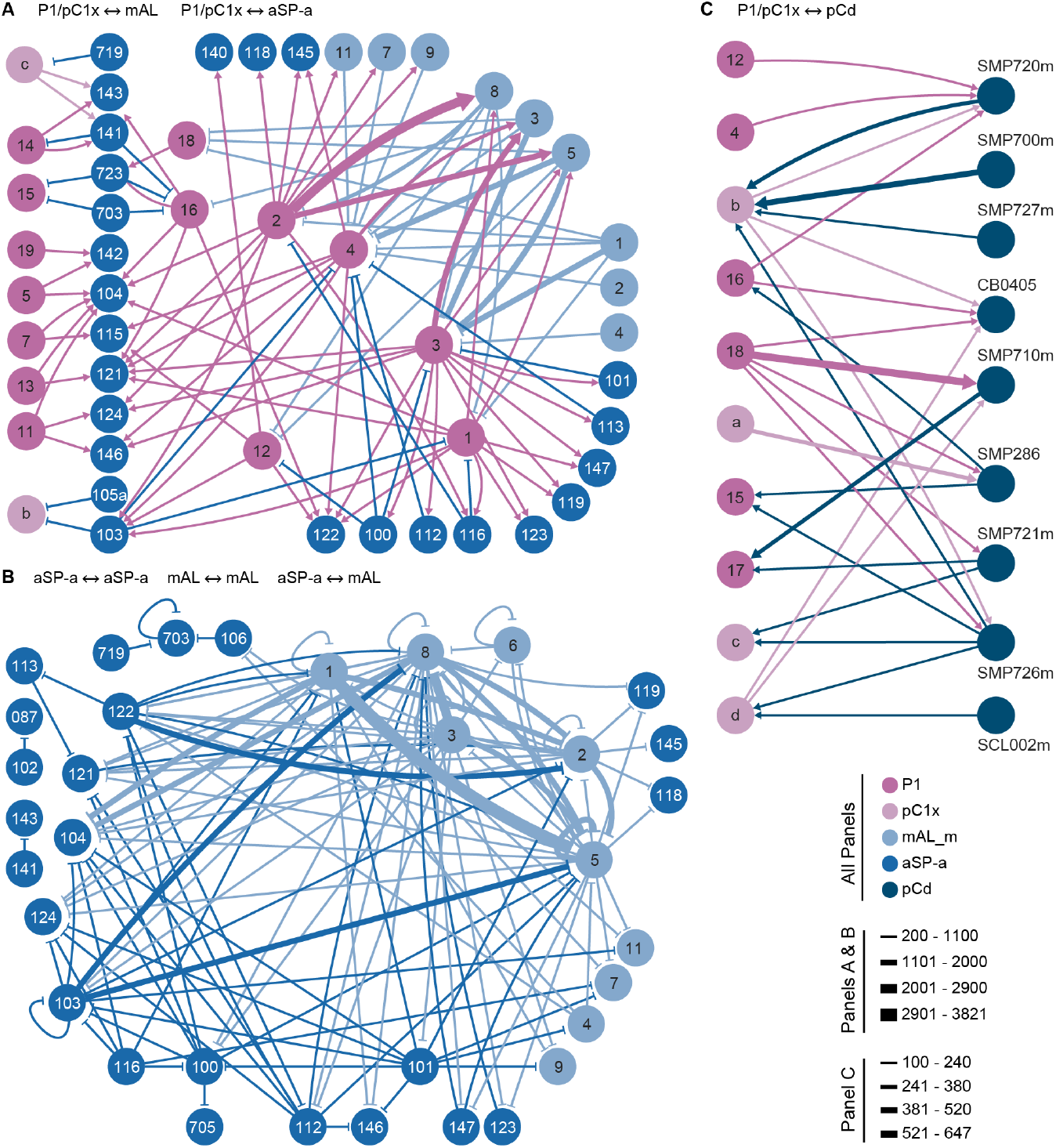
Diagrams of connections between P1, pC1x, aSP-a and mAL neurons. (A) Diagram of the same interactions shown in Figure 3C but with the threshold required to show a connection decreased from 500 to 200 synapses. (B) Diagram of the same interactions shown in Figure 3D but with the connection threshold decreased from 500 to 200 synapses. These diagrams give a more complete view of the networks connecting these cell types. The full names of the aSP-a cell types shown in these panels are: SIP100m, SIP101m, SIP103m, SIP104m, SIP 112m, SIP113m, SIP115m, SIP 116m, SIP 119m, SIP121m, SIP122m, SIP123m, SIP124m, SIP140m, SIP141m, SIP 142m, SIP 143m, SIP145m, SIP146m, SIP147m, SMP087, SMP088, SMP102, SMP105a, SMP106, SMP107, SMP703m, SMP705m, SMP719m, SMP723m. (C) Diagram of the connections between P1/cP1x supertypes and neurons from the pCd group of cell types, a group of sex-specific or sexually dimorphic cell types from the *dsx* -positive pCd lineage (51).

**Fig. S5.**
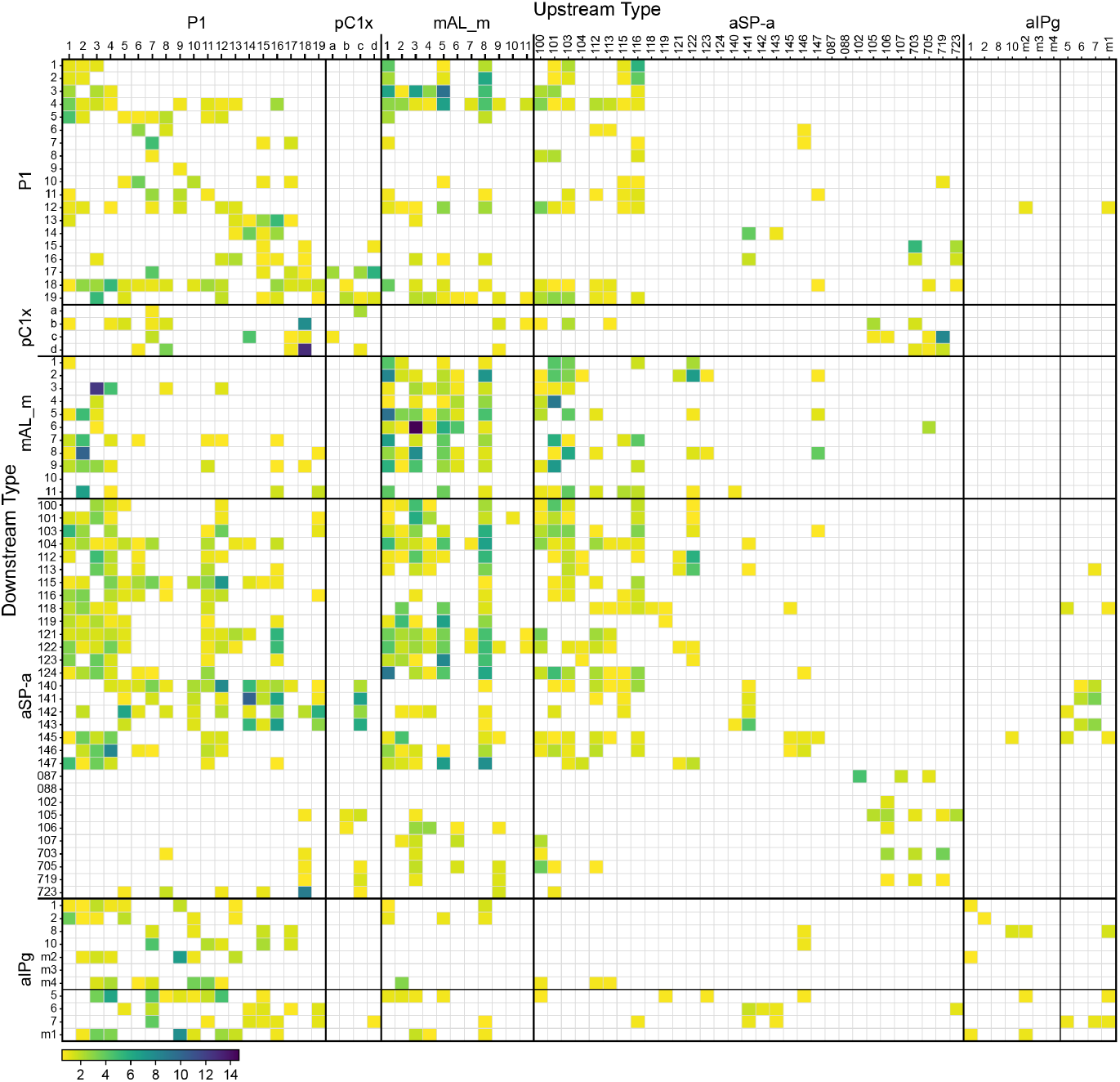
Connections between P1, pC1x, mAL aSP-a and aIPg cell types shown as a matrix. This matrix provides a view of the connections between the key sex-specific and dimorphic cell types discussed in this paper. Columns show input cell types and rows their downstream targets. The fill of each cell in the matrix represents, as a heat map, the percentage of the downstream cell type’s total synaptic input is provided by the upstream cell type. Only connections representing more than 0.5% of the downstream cell type’s synapses are shown. This matrix complements the circuit diagrams shown in Figure 3B-D and Figure S4 where connection strength is based purely on synapse number. The full names of the aSP-a cell types are: SIP100m, SIP101m, SIP103m, SIP104m, SIP 112m, SIP113m, SIP115m, SIP 116m, SIP 119m, SIP121m, SIP122m, SIP123m, SIP124m, SIP140m, SIP141m, SIP 142m, SIP 143m, SIP145m, SIP146m, SIP147m, SMP087, SMP088, SMP102, SMP105a, SMP106, SMP107, SMP703m, SMP705m, SMP719m, SMP723m. An enlarged matrix where individual P1 cell types, rather supertypes of cell types, is given in Supplemental File 1.

**Fig. S6.**
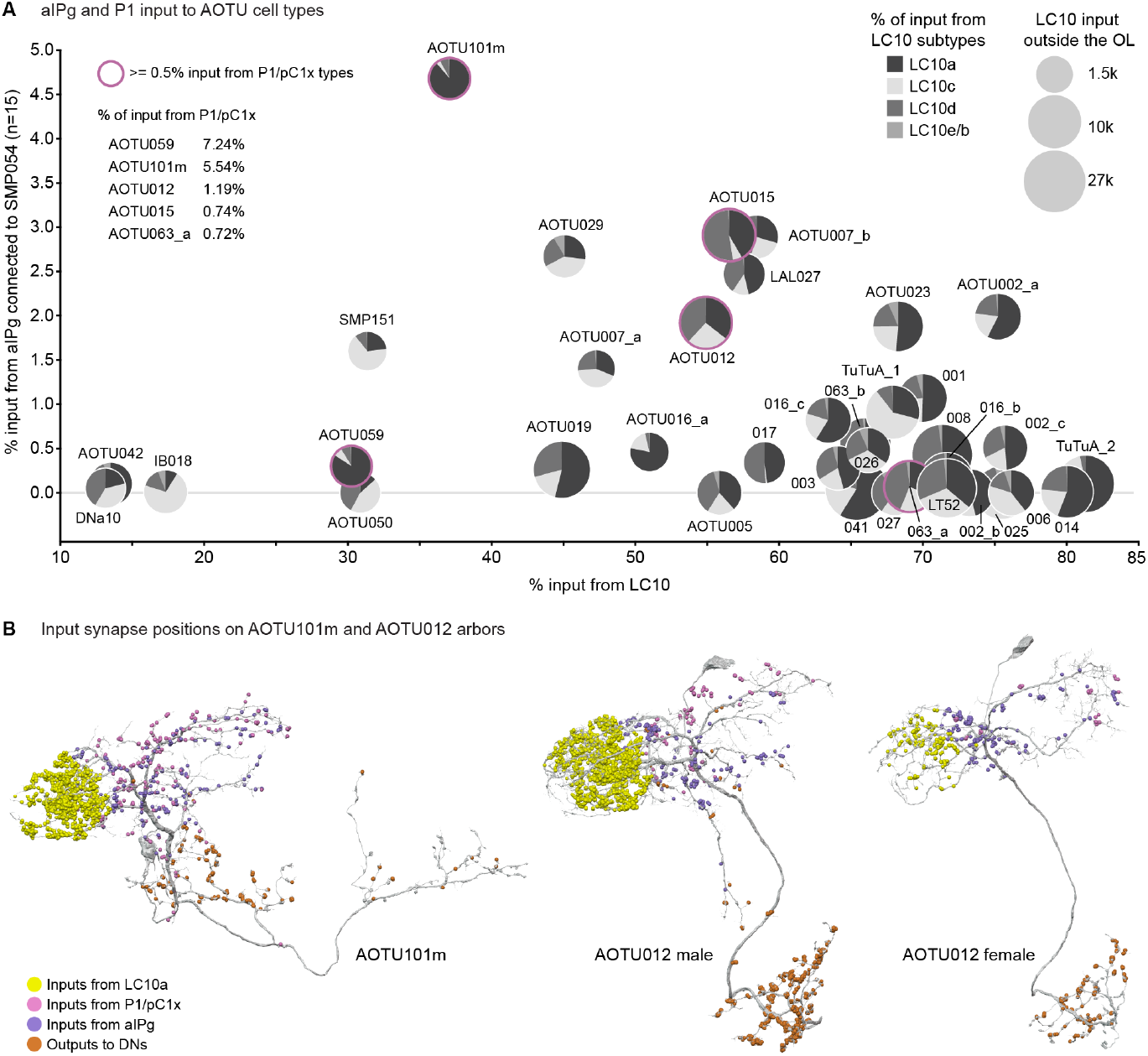
aIPg and P1 inputs to AOTU cell types. (A) AOTU cell types that receive input from LC10-group cell types and, in some cases, from aIPg or P1 cell types in the male. Each target cell type is represented by a circle whose diameter represents the total number of synapses it receives from LC10-group cell types. Some AOTU cell types are indicated as numbers without the “AOTU0” prefix. The proportion of those inputs coming from each LC10-group cell type is indicated as a pie chart. AOTU types that also get input from P1 neurons are outlined in purple. The percentage of input to those AOTU cell types provided by P1/pC1x neurons is shown. For comparison to the female, see Figure 2A in ref (3). (B) The positions of the color-coded input and output synapses on the arbors of AOTU101m, AOTU012 in males, and AOTU012 in females are shown.

**Fig. S7.**
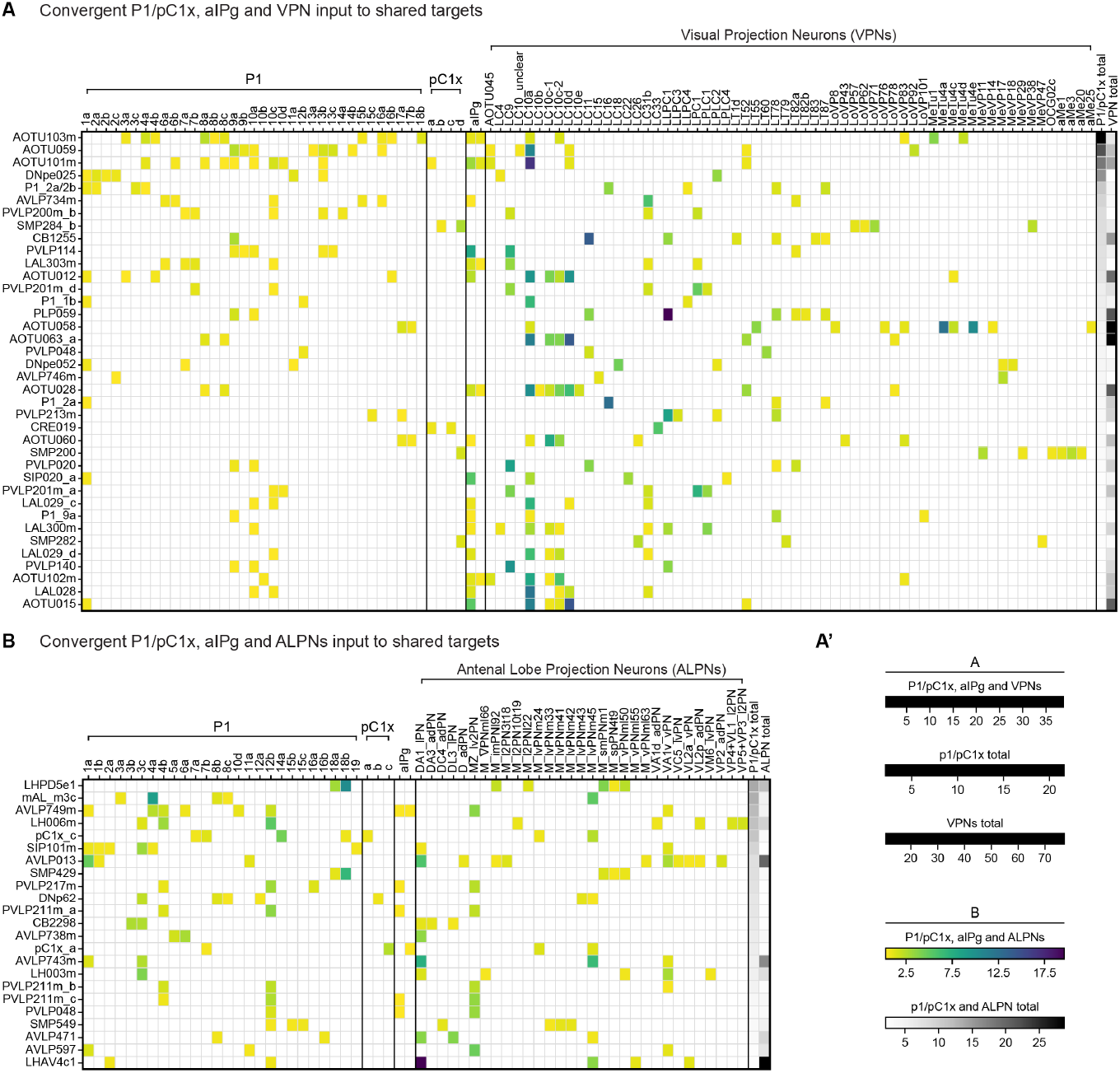
Matrices showing convergence of P1, pC1x, aIPg and sensory input neurons on common downstream targets. (A) Visual (VPN) sensory inputs. (B) Olfactory (ALPN) sensory inputs. Columns show input cell types and rows their downstream targets. The fill of each cell in the matrix represents, as heat maps (A’) the percentage of the downstream cell’s total synaptic input is provided by the upstream cell. Only connections representing more than 0.5% are shown.

## Video Supplements

**Video S1. Expression of fruitless, doublesex, ILP7 and DH44 in selected stable-split GAL4 lines**. A montage of z-stack videos showing of larger portions of the brains shown in Figure 2U-AL in which multiple cell bodies can be scored. Video available at: 10.25378/janelia.30392479.

**Video S2. Male-male behavior of upon activation of selected P1/pC1x cell types**. The expression pattern for each P1/pC1x split-GAL4 driver line is shown prior to the behavioral video for each line. Each video is displayed at 2x speed (120 FPS) and contains both a bird’s eye view of all nine arenas (Left) and a zoom into one arena from the same experiment (Right). The photoactivation (at 2.9 or 3.5 mW/cm^2^) period is indicated with a red dotted outline. Video available at: 10.25378/janelia.30400387.

**Video S3. Male group behavior upon activation of selected aIPg cell types**. The expression pattern for the aIPg split-GAL4 driver line is shown prior to the video. The same video (recorded at 120 FPS) is shown twice: first at 120 FPS and then slowed down at 35 FPS. The photoactivation (2.7 mW/cm^2^ at 660 nm) period is indicated with a red dotted outline. Video available at: 10.25378/janelia.30401053.

## Notes

https://doi.org/10.25378/janelia.30392479

https://doi.org/10.25378/janelia.30400387

https://doi.org/10.25378/janelia.30401053

### Competing Interest Statement

The authors have declared no competing interest.

## Bibliography

1. Catherine Dulac and Tali Kimchi. Neural mechanisms underlying sex-specific behaviors in vertebrates. Current Opinion in Neurobiology, 17(6):675–683, December 2007. ISSN 0959-4388. doi: 10.1016/j.conb.2008.01.009.

2. Meital Oren-Suissa and Troy R. Shirangi. Neural Circuits Underlying Sexually Dimorphic Innate Behaviors. Annual Review of Neuroscience, 48(1):191–210, July 2025. ISSN 0147-006X, 1545-4126. doi: 10.1146/annurev-neuro-112723-034621.

3. Catherine E. Schretter. Aggression across sexes from a contextual- and circuit-based perspective. Current Opinion in Neurobiology, 93:103071, August 2025. ISSN 0959-4388. doi: 10.1016/j.conb.2025.103071.

4. Cindy F. Yang and Nirao M. Shah. Representing sex in the brain, one module at a time. Neuron, 82(2):261–278, April 2014. ISSN 1097-4199. doi: 10.1016/j.neuron.2014.03.029.

5. Stephen F. Goodwin and Oliver Hobert. Molecular Mechanisms of Sexually Dimorphic Nervous System Patterning in Flies and Worms. Annual Review of Cell and Developmental Biology, 37(1):519–547, October 2021. ISSN 1081-0706, 1530-8995. doi: 10.1146/annurev-cellbio-120319-115237.

6. Thomas O Auer and Richard Benton. Sexual circuitry in Drosophila. Current Opinion in Neurobiology, 38:18–26, June 2016. ISSN 0959-4388. doi: 10.1016/j.conb.2016.01.004.

7. Kosei Sato and Daisuke Yamamoto. Molecular and cellular origins of behavioral sex differences: a tiny little fly tells a lot. Frontiers in Molecular Neuroscience, 16, October 2023. ISSN 1662-5099. doi: 10.3389/fnmol.2023.1284367. Publisher: Frontiers.

8. Gyunghee Lee, Margit Foss, Stephen F. Goodwin, Troy Carlo, Barbara J. Taylor, and Jeffrey C. Hall. Spatial, temporal, and sexually dimorphic expression patterns of the fruitless gene in the Drosophila central nervous system. Journal of Neurobiology, 43 (4):404–426, 2000. ISSN 1097-4695. doi: 10.1002/1097-4695(20000615)43:4<404::AID-NEU8>3.0.CO;2-D. _eprint: https://onlinelibrary.wiley.com/doi/pdf/10.1002/1097-4695%2820000615%2943%3A4%3C404%3A%3AAID-NEU8%3E3.0.CO%3B2-D.

9. Gyunghee Lee, Jeffrey C. Hall, and Jae H. Park. Doublesex Gene Expression in the Central Nervous System of Drosophila Melanogaster. Journal of Neurogenetics, 16(4): 229–248, January 2002. ISSN 0167-7063. doi: 10.1080/01677060216292. Publisher: Taylor & Francis _eprint: 10.1080/01677060216292.

10. Aaron M. Allen, Megan C. Neville, Tetsuya Nojima, Faredin Alejevski, and Stephen F. Goodwin. A Role for Exaptation in Sculpting Sexually Dimorphic Brains from Shared Neural Lineages, June 2025. Pages: 2025.06.04.657833 Section: New Results.

11. J M Belote and B S Baker. Sex determination in Drosophila melanogaster: analysis of transformer-2, a sex-transforming locus. Proceedings of the National Academy of Sciences, 79(5):1568–1572, March 1982. doi: 10.1073/pnas.79.5.1568. Publisher: Proceedings of the National Academy of Sciences.

12. B. S. Baker and M. F. Wolfner. A molecular analysis of doublesex, a bifunctional gene that controls both male and female sexual differentiation in Drosophila melanogaster. Genes & Development, 2(4):477–489, April 1988. ISSN 0890-9369. doi: 10.1101/gad.2.4.477.

13. Petra Stockinger, Duda Kvitsiani, Shay Rotkopf, László Tirián, and Barry J. Dickson. Neural Circuitry that Governs Drosophila Male Courtship Behavior. Cell, 121(5):795–807, June 2005. ISSN 0092-8674. doi: 10.1016/j.cell.2005.04.026.

14. Duda Kvitsiani and Barry J. Dickson. Shared neural circuitry for female and male sexual behaviours in Drosophila. Current Biology, 16(10):R355–R356, May 2006. ISSN 0960-9822. doi: 10.1016/j.cub.2006.04.025.

15. Elizabeth J. Rideout, Anthony J. Dornan, Megan C. Neville, Suzanne Eadie, and Stephen F. Goodwin. Control of sexual differentiation and behavior by the doublesex gene in Drosophila melanogaster. Nature Neuroscience, 13(4):458–466, April 2010. ISSN 1546-1726. doi: 10.1038/nn.2515. Publisher: Nature Publishing Group.

16. Troy R. Shirangi, Allan M. Wong, James W. Truman, and David L. Stern. Doublesex Regulates the Connectivity of a Neural Circuit Controlling Drosophila Male Courtship Song. Developmental Cell, 37(6):533–544, June 2016. ISSN 1534-5807. doi: 10.1016/j.devcel.2016.05.012.

17. Kenta Asahina. Sex differences in Drosophila behavior: qualitative and quantitative dimorphism. Current Opinion in Physiology, 6:35–45, December 2018. ISSN 2468-8673. doi: 10.1016/j.cophys.2018.04.004.

18. Stuart Berg, Isabella R. Beckett, Marta Costa, Philipp Schlegel, Michal Januszewski, Elizabeth C. Marin, Aljoscha Nern, Stephan Preibisch, Wei Qiu, Shin-ya Takemura, Alexandra M. C. Fragniere, Andrew S. Champion, Diane-Yayra Adjavon, Michael Cook, Marina Gkantia, Kenneth J. Hayworth, Gary B. Huang, Florian Kampf, William T. Katz, Zhiyuan Lu, Christopher Ordish, Tyler Paterson, Tomke Stuerner, Eric T. Trautman, Catherine R. Whittle, Laura E. Burnett, Judith Hoeller, Feng Li, Frank Loesche, Billy J. Morris, Tobias Pietzsch, Markus W. Pleijzier, Valeria Silva, Yijie Yin, Iris Ali, Griffin Badalamente, Alexander Shakeel Bates, John Bogovic, Paul Brooks, Sebastian Cachero, Brandon S. Canino, Bhumpanya Chaisrisawatsuk, Jody Clements, Arthur Crowe, Ines de Haan Vicente, Georgia Dempsey, Erika Dona, Marcia dos Santos, Marisa Dreher, Christopher R. Dunne, Katharina Eichler, Samantha Finley-May, Miriam A. Flynn, Imran Hameed, Gary Patrick Hopkins, Philip M. Hubbard, Ladann Kiassat, Julie Kovalyak, Shirley A. Lauchie, Meghan Leonard, Alanna Lohff, Kit D. Longden, Charli A. Maldonado, Myrto Mitletton, Ilina Moitra, Sung Soo Moon, Caroline Mooney, Eva J. Munnelly, Nneoma Okeoma, Donald J. Olbris, Anika Pai, Birava Patel, Emily M. Phillips, Stephen M. Plaza, Alana Richards, Jennifer Rivas Salinas, Ruairi J. V. Roberts, Edward M. Rogers, Ashley L. Scott, Louis A. Scuderi, Pavithraa Seenivasan, Laia Serratosa Capdevila, Claire Smith, Rob Svirskas, Satoko Takemura, Ibrahim Tastekin, Alexander Thomson, Lowell Umayam, John J. Walsh, Holly Whittome, C. Shan Xu, Emily A. Yakal, Tansy Yang, Arthur Zhao, Reed George, Viren Jain, Vivek Jayaraman, Wyatt Korff, Geoffrey W. Meissner, Sandro Romani, Jan Funke, Christopher Knecht, Stephan Saalfeld, Louis K. Scheffer, Scott Waddell, Gwyneth M. Card, Carlos Ribeiro, Michael B. Reiser, Harald F. Hess, Gerald M. Rubin, and Gregory S. X. E. Jefferis. Sexual dimorphism in the complete connectome of the Drosophila male central nervous system, October 2025. ISSN: 2692-8205 Pages: 2025.10.09.680999 Section: New Results.

19. Ken-Ichi Kimura, Tomoaki Hachiya, Masayuki Koganezawa, Tatsunori Tazawa, and Daisuke Yamamoto. Fruitless and doublesex coordinate to generate male-specific neurons that can initiate courtship. Neuron, 59(5):759–769, September 2008. ISSN 1097-4199. doi: 10.1016/j.neuron.2008.06.007.

20. Qingzhong Ren, Takeshi Awasaki, Yu-Fen Huang, Zhiyong Liu, and Tzumin Lee. Cell Class-Lineage Analysis Reveals Sexually Dimorphic Lineage Compositions in the Drosophila Brain. Current Biology, 26(19):2583–2593, October 2016. ISSN 0960-9822. doi: 10.1016/j.cub.2016.07.086.

21. Soh Kohatsu, Masayuki Koganezawa, and Daisuke Yamamoto. Female Contact Activates Male-Specific Interneurons that Trigger Stereotypic Courtship Behavior in Drosophila. Neuron, 69(3):498–508, February 2011. ISSN 0896-6273. doi: 10.1016/j.neuron.2010.12.017.

22. Anne C. von Philipsborn, Tianxiao Liu, Jai Y. Yu, Christopher Masser, Salil S. Bidaye, and Barry J. Dickson. Neuronal Control of Drosophila Courtship Song. Neuron, 69(3):509–522, February 2011. ISSN 0896-6273. doi: 10.1016/j.neuron.2011.01.011.

23. Yufeng Pan, Geoffrey W. Meissner, and Bruce S. Baker. Joint control of Drosophila male courtship behavior by motion cues and activation of male-specific P1 neurons. Proceedings of the National Academy of Sciences, 109(25):10065–10070, June 2012. doi: 10.1073/pnas.1207107109. Publisher: Proceedings of the National Academy of Sciences.

24. Soh Kohatsu and Daisuke Yamamoto. Visually induced initiation of Drosophila innate courtship-like following pursuit is mediated by central excitatory state. Nature Communications, 6(1):6457, March 2015. ISSN 2041-1723. doi: 10.1038/ncomms7457. Publisher: Nature Publishing Group.

25. E. Josephine Clowney, Shinya Iguchi, Jennifer J. Bussell, Elias Scheer, and Vanessa Ruta. Multimodal Chemosensory Circuits Controlling Male Courtship in Drosophila. Neuron, 87 (5):1036–1049, September 2015. ISSN 0896-6273. doi: 10.1016/j.neuron.2015.07.025.

26. István Taisz, Erika Donà, Daniel Münch, Shanice N. Bailey, Billy J. Morris, Kimberly I. Meechan, Katie M. Stevens, Irene Varela-Martínez, Marina Gkantia, Philipp Schlegel, Carlos Ribeiro, Gregory S. X. E. Jefferis, and Dana S. Galili. Generating parallel representations of position and identity in the olfactory system. Cell, 186(12):2556–2573.e22, June 2023. ISSN 0092-8674. doi: 10.1016/j.cell.2023.04.038.

27. Sweta Agrawal and Michael H. Dickinson. The effects of target contrast on Drosophila courtship. The Journal of Experimental Biology, 222(Pt 16):jeb203414, August 2019. ISSN 1477-9145. doi: 10.1242/jeb.203414.

28. Benjamin R Kallman, Heesoo Kim, and Kristin Scott. Excitation and inhibition onto central courtship neurons biases Drosophila mate choice. eLife, 4:e11188, November 2015. ISSN 2050-084X. doi: 10.7554/eLife.11188. Publisher: eLife Sciences Publications, Ltd.

29. Dandan Chen, Divya Sitaraman, Nan Chen, Xin Jin, Caihong Han, Jie Chen, Mengshi Sun, Bruce S. Baker, Michael N. Nitabach, and Yufeng Pan. Genetic and neuronal mechanisms governing the sex-specific interaction between sleep and sexual behaviors in Drosophila. Nature Communications, 8(1):154, July 2017. ISSN 2041-1723. doi: 10.1038/s41467-017-00087-5. Publisher: Nature Publishing Group.

30. Masayuki Koganezawa, Ken-ichi Kimura, and Daisuke Yamamoto. The Neural Circuitry that Functions as a Switch for Courtship versus Aggression in Drosophila Males. Current Biology, 26(11):1395–1403, June 2016. ISSN 0960-9822. doi: 10.1016/j.cub.2016.04.017.

31. Eric D Hoopfer, Yonil Jung, Hidehiko K Inagaki, Gerald M Rubin, and David J Anderson. P1 interneurons promote a persistent internal state that enhances inter-male aggression in Drosophila. eLife, 4:e11346, December 2015. ISSN 2050-084X. doi: 10.7554/eLife.11346. Publisher: eLife Sciences Publications, Ltd.

32. Kiichi Watanabe, Hui Chiu, and David J Anderson. Whole-brain in situ mapping of neuronal activation in Drosophila during social behaviors and optogenetic stimulation. eLife, 12: RP92380, November 2024. ISSN 2050-084X. doi: 10.7554/eLife.92380. Publisher: eLife Sciences Publications, Ltd.

33. Tom Hindmarsh Sten, Rufei Li, Florian Hollunder, Shade Eleazer, and Vanessa Ruta. Male-male interactions shape mate selection in Drosophila. Cell, 188(6):1486–1503.e25, March 2025. ISSN 0092-8674. doi: 10.1016/j.cell.2025.01.008.

34. Daniel E. Bath, John R. Stowers, Dorothea Hörmann, Andreas Poehlmann, Barry J. Dickson, and Andrew D. Straw. FlyMAD: rapid thermogenetic control of neuronal activity in freely walking Drosophila. Nature Methods, 11(7):756–762, July 2014. ISSN 1548-7105. doi: 10.1038/nmeth.2973. Publisher: Nature Publishing Group.

35. Hidehiko K. Inagaki, Yonil Jung, Eric D. Hoopfer, Allan M. Wong, Neeli Mishra, John Y. Lin, Roger Y. Tsien, and David J. Anderson. Optogenetic control of Drosophila using a redshifted channelrhodopsin reveals experience-dependent influences on courtship. Nature Methods, 11(3):325–332, March 2014. ISSN 1548-7105. doi: 10.1038/nmeth.2765.

36. Chuan Zhou, Yufeng Pan, Carmen C. Robinett, Geoffrey W. Meissner, and Bruce S. Baker. Central Brain Neurons Expressing doublesex Regulate Female Receptivity in Drosophila. Neuron, 83(1):149–163, July 2014. ISSN 0896-6273. doi: 10.1016/j.neuron.2014.05.038.

37. Kaiyu Wang, Fei Wang, Nora Forknall, Tansy Yang, Christopher Patrick, Ruchi Parekh, and Barry J. Dickson. Neural circuit mechanisms of sexual receptivity in Drosophila females. Nature, 589(7843):577–581, January 2021. ISSN 1476-4687. doi: 10.1038/s41586-020-2972-7. Publisher: Nature Publishing Group.

38. David Deutsch, Diego Pacheco, Lucas Encarnacion-Rivera, Talmo Pereira, Ramie Fathy, Jan Clemens, Cyrille Girardin, Adam Calhoun, Elise Ireland, Austin Burke, Sven Dorkenwald, Claire McKellar, Thomas Macrina, Ran Lu, Kisuk Lee, Nico Kemnitz, Dodam Ih, Manuel Castro, Akhilesh Halageri, Chris Jordan, William Silversmith, Jingpeng Wu, H Sebastian Seung, and Mala Murthy. The neural basis for a persistent internal state in Drosophila females. eLife, 9:e59502, November 2020. ISSN 2050-084X. doi: 10.7554/eLife.59502. Publisher: eLife Sciences Publications, Ltd.

39. Catherine E Schretter, Yoshinori Aso, Alice A Robie, Marisa Dreher, Michael-John Dolan, Nan Chen, Masayoshi Ito, Tansy Yang, Ruchi Parekh, Kristin M Branson, and Gerald M Rubin. Cell types and neuronal circuitry underlying female aggression in Drosophila. eLife, 9:e58942, November 2020. ISSN 2050-084X. doi: 10.7554/eLife.58942. Publisher: eLife Sciences Publications, Ltd.

40. Hui Chiu, Eric D. Hoopfer, Maeve L. Coughlan, Hania J. Pavlou, Stephen F. Goodwin, and David J. Anderson. A circuit logic for sexually shared and dimorphic aggressive behaviors in Drosophila. Cell, 184(3):847, February 2021. ISSN 0092-8674. doi: 10.1016/j.cell.2021.01.021.

41. Hui Chiu, Alice A Robie, Kristin Branson, Tanvi Vippa, Samantha Epstein, Gerald M Rubin, David J Anderson, and Catherine E Schretter. Cell type-specific contributions to a persistent aggressive internal state in female Drosophila. eLife, 12:RP88598, July 2025. ISSN 2050-084X. doi: 10.7554/eLife.88598. Publisher: eLife Sciences Publications, Ltd.

42. Fei Wang, Kaiyu Wang, Nora Forknall, Christopher Patrick, Tansy Yang, Ruchi Parekh, Davi Bock, and Barry J. Dickson. Neural circuitry linking mating and egg laying in Drosophila females. Nature, 579(7797):101–105, March 2020. ISSN 1476-4687. doi: 10.1038/s41586-020-2055-9. Publisher: Nature Publishing Group.

43. Caroline B. Palavicino-Maggio, Yick-Bun Chan, Claire McKellar, and Edward A. Kravitz. A small number of cholinergic neurons mediate hyperaggression in female Drosophila. Proceedings of the National Academy of Sciences, 116(34):17029–17038, August 2019. doi: 10.1073/pnas.1907042116. Publisher: Proceedings of the National Academy of Sciences.

44. Sebastian Cachero, Aaron D. Ostrovsky, Jai Y. Yu, Barry J. Dickson, and Gregory S. X. E. Jefferis. Sexual Dimorphism in the Fly Brain. Current Biology, 20(18):1589–1601, September 2010. ISSN 0960-9822. doi: 10.1016/j.cub.2010.07.045.

45. Louis K. Scheffer, C. Shan Xu, Michal Januszewski, Zhiyuan Lu, Shin-Ya Takemura, Kenneth J. Hayworth, Gary B. Huang, Kazunori Shinomiya, Jeremy Maitlin-Shepard, Stuart Berg, Jody Clements, Philip M. Hubbard, William T. Katz, Lowell Umayam, Ting Zhao, David Ackerman, Tim Blakely, John Bogovic, Tom Dolafi, Dagmar Kainmueller, Takashi Kawase, Khaled A. Khairy, Laramie Leavitt, Peter H. Li, Larry Lindsey, Nicole Neubarth, Donald J. Olbris, Hideo Otsuna, Eric T. Trautman, Masayoshi Ito, Alexander S. Bates, Jens Goldammer, Tanya Wolff, Robert Svirskas, Philipp Schlegel, Erika Neace, Christopher J. Knecht, Chelsea X. Alvarado, Dennis A. Bailey, Samantha Ballinger, Jolanta A. Borycz, Brandon S. Canino, Natasha Cheatham, Michael Cook, Marisa Dreher, Octave Duclos, Bryon Eubanks, Kelli Fairbanks, Samantha Finley, Nora Forknall, Audrey Francis, Gary Patrick Hopkins, Emily M. Joyce, SungJin Kim, Nicole A. Kirk, Julie Kovalyak, Shirley A. Lauchie, Alanna Lohff, Charli Maldonado, Emily A. Manley, Sari McLin, Caroline Mooney, Miatta Ndama, Omotara Ogundeyi, Nneoma Okeoma, Christopher Ordish, Nicholas Padilla, Christopher M. Patrick, Tyler Paterson, Elliott E. Phillips, Emily M. Phillips, Neha Rampally, Caitlin Ribeiro, Madelaine K. Robertson, Jon Thomson Rymer, Sean M. Ryan, Megan Sammons, Anne K. Scott, Ashley L. Scott, Aya Shinomiya, Claire Smith, Kelsey Smith, Natalie L. Smith, Margaret A. Sobeski, Alia Suleiman, Jackie Swift, Satoko Takemura, Iris Talebi, Dorota Tarnogorska, Emily Tenshaw, Temour Tokhi, John J. Walsh, Tansy Yang, Jane Anne Horne, Feng Li, Ruchi Parekh, Patricia K. Rivlin, Vivek Jayaraman, Marta Costa, Gregory Sxe Jefferis, Kei Ito, Stephan Saalfeld, Reed George, Ian A. Meinertzhagen, Gerald M. Rubin, Harald F. Hess, Viren Jain, and Stephen M. Plaza. A connectome and analysis of the adult Drosophila central brain. eLife, 9:e57443, September 2020. ISSN 2050-084X. doi: 10.7554/eLife.57443.

46. Sven Dorkenwald, Arie Matsliah, Amy R. Sterling, Philipp Schlegel, Szi-chieh Yu, Claire E. McKellar, Albert Lin, Marta Costa, Katharina Eichler, Yijie Yin, Will Silversmith, Casey Schneider-Mizell, Chris S. Jordan, Derrick Brittain, Akhilesh Halageri, Kai Kuehner, Oluwaseun Ogedengbe, Ryan Morey, Jay Gager, Krzysztof Kruk, Eric Perlman, Runzhe Yang, David Deutsch, Doug Bland, Marissa Sorek, Ran Lu, Thomas Macrina, Kisuk Lee, J. Alexander Bae, Shang Mu, Barak Nehoran, Eric Mitchell, Sergiy Popovych, Jingpeng Wu, Zhen Jia, Manuel A. Castro, Nico Kemnitz, Dodam Ih, Alexander Shakeel Bates, Nils Eckstein, Jan Funke, Forrest Collman, Davi D. Bock, Gregory S. X. E. Jefferis, H. Sebastian Seung, and Mala Murthy. Neuronal wiring diagram of an adult brain. Nature, 634(8032):124–138, October 2024. ISSN 1476-4687. doi: 10.1038/s41586-024-07558-y. Publisher: Nature Publishing Group.

47. Philipp Schlegel, Yijie Yin, Alexander S. Bates, Sven Dorkenwald, Katharina Eichler, Paul Brooks, Daniel S. Han, Marina Gkantia, Marcia dos Santos, Eva J. Munnelly, Griffin Badalamente, Laia Serratosa Capdevila, Varun A. Sane, Alexandra M. C. Fragniere, Ladann Kiassat, Markus W. Pleijzier, Tomke Stürner, Imaan F. M. Tamimi, Christopher R. Dunne, Irene Salgarella, Alexandre Javier, Siqi Fang, Eric Perlman, Tom Kazimiers, Sridhar R. Jagannathan, Arie Matsliah, Amy R. Sterling, Szi-chieh Yu, Claire E. McKellar, Marta Costa, H. Sebastian Seung, Mala Murthy, Volker Hartenstein, Davi D. Bock, and Gregory S. X. E. Jefferis. Whole-brain annotation and multi-connectome cell typing of Drosophila. Nature, 634(8032):139–152, October 2024. ISSN 1476-4687. doi: 10.1038/s41586-024-07686-5. Publisher: Nature Publishing Group.

48. Zhihao Zheng, J. Scott Lauritzen, Eric Perlman, Camenzind G. Robinson, Matthew Nichols, Daniel Milkie, Omar Torrens, John Price, Corey B. Fisher, Nadiya Sharifi, Steven A. Calle-Schuler, Lucia Kmecova, Iqbal J. Ali, Bill Karsh, Eric T. Trautman, John A. Bogovic, Philipp Hanslovsky, Gregory S. X. E. Jefferis, Michael Kazhdan, Khaled Khairy, Stephan Saalfeld, Richard D. Fetter, and Davi D. Bock. A Complete Electron Microscopy Volume of the Brain of Adult Drosophila melanogaster. Cell, 174(3):730–743.e22, July 2018. ISSN 0092-8674. doi: 10.1016/j.cell.2018.06.019.

49. David Deutsch, Arie Matsliah, Kaiyu Wang, Sven Dorkenwald, Arpita Mondal, Austin Burke, James Hebditch, Jay Gager, Szi-Chieh Yu, Amy Sterling, Claire McKellar, Philipp Schlegel, Stephan Gerhard, Gabriella Sterne, Marta Costa, Katharina Eichler, Yijie Yin, S. X. E. Gregory, Barry Dickson, H. Sebastian Seung, and Mala Murthy. Sexually-dimorphic neurons in the Drosophila whole-brain connectome. Research Square, pages rs.3.rs–6881911, June 2025. ISSN 2693-5015. doi: 10.21203/rs.3.rs-6881911/v1.

50. Masayuki Koganezawa, Daisuke Haba, Takashi Matsuo, and Daisuke Yamamoto. The Shaping of Male Courtship Posture by Lateralized Gustatory Inputs to Male-Specific Interneurons. Current Biology, 20(1):1–8, January 2010. ISSN 0960-9822. doi: 10.1016/j.cub.2009.11.038.

51. Tetsuya Nojima, Annika Rings, Aaron M. Allen, Nils Otto, Thomas A. Verschut, Jean-Christophe Billeter, Megan C. Neville, and Stephen F. Goodwin. A sex-specific switch between visual and olfactory inputs underlies adaptive sex differences in behavior. Current Biology, 31(6):1175–1191.e6, March 2021. ISSN 0960-9822. doi: 10.1016/j.cub.2020.12.047.

52. Ken-Ichi Kimura, Manabu Ote, Tatsunori Tazawa, and Daisuke Yamamoto. Fruitless specifies sexually dimorphic neural circuitry in the Drosophila brain. Nature, 438(7065):229–233, November 2005. ISSN 1476-4687. doi: 10.1038/nature04229.

53. Joshua L. Lillvis, Kaiyu Wang, Hiroshi M. Shiozaki, Min Xu, David L. Stern, and Barry J. Dickson. Nested neural circuits generate distinct acoustic signals during Drosophila courtship. Current biology: CB, 34(4):808–824.e6, February 2024. ISSN 1879-0445. doi: 10.1016/j.cub.2024.01.015.

54. Heather Dionne, Karen L. Hibbard, Amanda Cavallaro, Jui-Chun Kao, and Gerald M. Rubin. Genetic Reagents for Making Split-GAL4 Lines in Drosophila. Genetics, 209(1):31–35, May 2018. ISSN 1943-2631. doi: 10.1534/genetics.118.300682.

55. Geoffrey W. Meissner, Aljoscha Nern, Zachary Dorman, Gina M. DePasquale, Kaitlyn Forster, Theresa Gibney, Joanna H. Hausenfluck, Yisheng He, Nirmala A. Iyer, Jennifer Jeter, Lauren Johnson, Rebecca M. Johnston, Kelley Lee, Brian Melton, Brianna Yarbrough, Christopher T. Zugates, Jody Clements, Cristian Goina, Hideo Otsuna, Konrad Rokicki, Robert R. Svirskas, Yoshinori Aso, Gwyneth M. Card, Barry J. Dickson, Erica Ehrhardt, Jens Goldammer, Masayoshi Ito, Dagmar Kainmueller, Wyatt Korff, Lisa Mais, Ryo Minegishi, Shigehiro Namiki, Gerald M. Rubin, Gabriella R. Sterne, Tanya Wolff, Oz Malkesman, and FlyLight Project Team. A searchable image resource of Drosophila GAL4 driver expression patterns with single neuron resolution. eLife, 12:e80660, February 2023. ISSN 2050-084X. doi: 10.7554/eLife.80660.

56. Laszlo Tirian and Barry J. Dickson. The VT GAL4, LexA, and split-GAL4 driver line collections for targeted expression in the Drosophila nervous system, October 2017. Pages: 198648 Section: New Results.

57. Aljoscha Nern, Frank Loesche, Shin-Ya Takemura, Laura E. Burnett, Marisa Dreher, Eyal Gruntman, Judith Hoeller, Gary B. Huang, Michał Januszewski, Nathan C. Klapoetke, Sanna Koskela, Kit D. Longden, Zhiyuan Lu, Stephan Preibisch, Wei Qiu, Edward M. Rogers, Pavithraa Seenivasan, Arthur Zhao, John Bogovic, Brandon S. Canino, Jody Clements, Michael Cook, Samantha Finley-May, Miriam A. Flynn, Imran Hameed, Alexandra M. C. Fragniere, Kenneth J. Hayworth, Gary Patrick Hopkins, Philip M. Hubbard, William T. Katz, Julie Kovalyak, Shirley A. Lauchie, Meghan Leonard, Alanna Lohff, Charli A. Maldonado, Caroline Mooney, Nneoma Okeoma, Donald J. Olbris, Christopher Ordish, Tyler Paterson, Emily M. Phillips, Tobias Pietzsch, Jennifer Rivas Salinas, Patricia K. Rivlin, Philipp Schlegel, Ashley L. Scott, Louis A. Scuderi, Satoko Takemura, Iris Talebi, Alexander Thomson, Eric T. Trautman, Lowell Umayam, Claire Walsh, John J. Walsh, C. Shan Xu, Emily A. Yakal, Tansy Yang, Ting Zhao, Jan Funke, Reed George, Harald F. Hess, Gregory S. X. E. Jefferis, Christopher Knecht, Wyatt Korff, Stephen M. Plaza, Sandro Romani, Stephan Saalfeld, Louis K. Scheffer, Stuart Berg, Gerald M. Rubin, and Michael B. Reiser. Connectome-driven neural inventory of a complete visual system. Nature, 641(8065):1225–1237, May 2025. ISSN 1476-4687. doi: 10.1038/s41586-025-08746-0.

58. Daisuke Yamamoto and Masayuki Koganezawa. Genes and circuits of courtship behaviour in Drosophila males. Nature Reviews Neuroscience, 14(10):681–692, October 2013. ISSN 1471-0048. doi: 10.1038/nrn3567. Publisher: Nature Publishing Group.

59. Osama M. Ahmed, Amanda Crocker, and Mala Murthy. Transcriptional profiling of Drosophila male-specific P1 (pC1) neurons, November 2023. Pages: 2023.11.07.566045 Section: New Results.

60. Jai Y. Yu, Makoto I. Kanai, Ebru Demir, Gregory S. X. E. Jefferis, and Barry J. Dickson. Cellular Organization of the Neural Circuit that Drives Drosophila Courtship Behavior. Current Biology, 20(18):1602–1614, September 2010. ISSN 0960-9822. doi: 10.1016/j.cub.2010.08.025.

61. Kiichi Watanabe, Hui Chiu, Barret D. Pfeiffer, Allan M. Wong, Eric D. Hoopfer, Gerald M. Rubin, and David J. Anderson. A Circuit Node that Integrates Convergent Input from Neuromodulatory and Social Behavior-Promoting Neurons to Control Aggression in Drosophila. Neuron, 95(5):1112–1128.e7, August 2017. ISSN 0896-6273. doi: 10.1016/j.neuron.2017.08.017.

62. Yonil Jung, Ann Kennedy, Hui Chiu, Farhan Mohammad, Adam Claridge-Chang, and David J. Anderson. Neurons that Function within an Integrator to Promote a Persistent Behavioral State in Drosophila. Neuron, 105(2):322–333.e5, January 2020. ISSN 1097-4199. doi: 10.1016/j.neuron.2019.10.028.

63. Inês M. A. Ribeiro, Michael Drews, Armin Bahl, Christian Machacek, Alexander Borst, and Barry J. Dickson. Visual Projection Neurons Mediating Directed Courtship in Drosophila. Cell, 174(3):607–621.e18, July 2018. ISSN 0092-8674, 1097-4172. doi: 10.1016/j.cell.2018.06.020.

64. Tom Hindmarsh Sten, Rufei Li, Adriane Otopalik, and Vanessa Ruta. Sexual arousal gates visual processing during Drosophila courtship. Nature, 595(7868):549–553, July 2021. ISSN 1476-4687. doi: 10.1038/s41586-021-03714-w.

65. Ming Wu, Aljoscha Nern, W. Ryan Williamson, Mai M. Morimoto, Michael B. Reiser, Gwyneth M. Card, and Gerald M. Rubin. Visual projection neurons in the Drosophila lobula link feature detection to distinct behavioral programs. eLife, 5:e21022, December 2016. ISSN 2050-084X. doi: 10.7554/eLife.21022.

66. Rajyashree Sen, Ming Wu, Kristin Branson, Alice Robie, Gerald M. Rubin, and Barry J. Dickson. Moonwalker Descending Neurons Mediate Visually Evoked Retreat in Drosophila. Current Biology, 27(5):766–771, March 2017. ISSN 0960-9822. doi: 10.1016/j.cub.2017.02.008.

67. Vinoy Vijayan, Rob Thistle, Tong Liu, Elena Starostina, and Claudio W. Pikielny. Drosophila pheromone-sensing neurons expressing the ppk25 ion channel subunit stimulate male courtship and female receptivity. PLoS genetics, 10(3):e1004238, March 2014. ISSN 1553-7404. doi: 10.1371/journal.pgen.1004238.

68. Amina Kurtovic, Alexandre Widmer, and Barry J. Dickson. A single class of olfactory neurons mediates behavioural responses to a Drosophila sex pheromone. Nature, 446 (7135):542–546, March 2007. ISSN 1476-4687. doi: 10.1038/nature05672. Publisher: Nature Publishing Group.

69. Aki Ejima, Benjamin P.C. Smith, Christophe Lucas, Wynand Van der Goes van Naters, Carson J. Miller, John R. Carlson, Joel D. Levine, and Leslie C. Griffith. Generalization of courtship learning in Drosophila is mediated by cis-vaccenyl acetate. Current biology : CB, 17(7):599–605, April 2007. ISSN 0960-9822. doi: 10.1016/j.cub.2007.01.053.

70. Weiwei Liu, Xinhua Liang, Jianxian Gong, Zhen Yang, Yao-Hua Zhang, Jian-Xu Zhang, and Yi Rao. Social regulation of aggression by pheromonal activation of Or65a olfactory neurons in Drosophila. Nature Neuroscience, 14(7):896–902, June 2011. ISSN 1546-1726. doi: 10.1038/nn.2836.

71. Liming Wang and David J. Anderson. Identification of an aggression-promoting pheromone and its receptor neurons in Drosophila. Nature, 463(7278):227–231, January 2010. ISSN 1476-4687. doi: 10.1038/nature08678. Publisher: Nature Publishing Group.

72. Vanessa Ruta, Sandeep Robert Datta, Maria Luisa Vasconcelos, Jessica Freeland, Loren L. Looger, and Richard Axel. A dimorphic pheromone circuit in Drosophila from sensory input to descending output. Nature, 468(7324):686–690, December 2010. ISSN 1476-4687. doi: 10.1038/nature09554. Publisher: Nature Publishing Group.

73. Kenta Asahina, Kiichi Watanabe, Brian J. Duistermars, Eric Hoopfer, Carlos Roberto González, Eyrún Arna Eyjólfsdóttir, Pietro Perona, and David J. Anderson. Tachykinin-expressing neurons control male-specific aggressive arousal in Drosophila. Cell, 156(0): 221–235, January 2014. ISSN 0092-8674. doi: 10.1016/j.cell.2013.11.045.

74. Johannes Kohl, Aaron D. Ostrovsky, Shahar Frechter, and Gregory S.X.E. Jefferis. A Bidirectional Circuit Switch Reroutes Pheromone Signals in Male and Female Brains. Cell, 155(7):1610–1623, December 2013. ISSN 0092-8674. doi: 10.1016/j.cell.2013.11.025.

75. Isabella R. Beckett, William Morris, and et al. Connectomic analysis of sexually dimorphic pheromone pathways in male Drosophila. in prep.

76. Julie H. Simpson. Descending control of motor sequences in Drosophila. Current Opinion in Neurobiology, 84:102822, February 2024. ISSN 0959-4388. doi: 10.1016/j.conb.2023.102822.

77. Tomke Stürner, Paul Brooks, Laia Serratosa Capdevila, Billy J. Morris, Alexandre Javier, Siqi Fang, Marina Gkantia, Sebastian Cachero, Isabella R. Beckett, Elizabeth C. Marin, Philipp Schlegel, Andrew S. Champion, Ilina Moitra, Alana Richards, Finja Klemm, Leonie Kugel, Shigehiro Namiki, Han S. J. Cheong, Julie Kovalyak, Emily Tenshaw, Ruchi Parekh, Jasper S. Phelps, Brandon Mark, Sven Dorkenwald, Alexander S. Bates, Arie Matsliah, Szi-chieh Yu, Claire E. McKellar, Amy Sterling, H. Sebastian Seung, Mala Murthy, John C. Tuthill, Wei-Chung Allen Lee, Gwyneth M. Card, Marta Costa, Gregory S. X. E. Jefferis, and Katharina Eichler. Comparative connectomics of Drosophila descending and ascending neurons. Nature, 643(8070):158–172, July 2025. ISSN 1476-4687. doi: 10.1038/s41586-025-08925-z. Publisher: Nature Publishing Group.

78. Shigehiro Namiki, Michael H. Dickinson, Allan M. Wong, Wyatt Korff, and Gwyneth M. Card. The functional organization of descending sensory-motor pathways in Drosophila. eLife, 7:e34272, June 2018. ISSN 2050-084X. doi: 10.7554/eLife.34272.

79. Claire E. McKellar, Joshua L. Lillvis, Daniel E. Bath, James E. Fitzgerald, John G. Cannon, Julie H. Simpson, and Barry J. Dickson. Threshold-Based Ordering of Sequential Actions during Drosophila Courtship. Current Biology, 29(3):426–434.e6, February 2019. ISSN 0960-9822. doi: 10.1016/j.cub.2018.12.019.

80. Frederic A. Roemschied, Diego A. Pacheco, Max J. Aragon, Elise C. Ireland, Xinping Li, Kyle Thieringer, Rich Pang, and Mala Murthy. Flexible circuit mechanisms for contextdependent song sequencing. Nature, 622(7984):794–801, October 2023. ISSN 1476-4687. doi: 10.1038/s41586-023-06632-1. Publisher: Nature Publishing Group.

81. Nathan C. Klapoetke, Yasunobu Murata, Sung Soo Kim, Stefan R. Pulver, Amanda Birdsey-Benson, Yong Ku Cho, Tania K. Morimoto, Amy S. Chuong, Eric J. Carpenter, Zhijian Tian, Jun Wang, Yinlong Xie, Zhixiang Yan, Yong Zhang, Brian Y. Chow, Barbara Surek, Michael Melkonian, Vivek Jayaraman, Martha Constantine-Paton, Gane Ka-Shu Wong, and Edward S. Boyden. Independent optical excitation of distinct neural populations. Nature Methods, 11(3):338–346, March 2014. ISSN 1548-7105. doi: 10.1038/nmeth.2836.

82. Steve Sawtelle, Lakshmi Narayan, Yun Ding, Elizabeth Kim, Emily L. Behrman, Joshua L. Lillvis, Takashi Kawase, and David L. Stern. Song Torrent: A modular, open-source 96-chamber audio and video recording apparatus with optogenetic activation and inactivation capabilities for Drosophila, January 2024. Pages: 2024.01.09.574712 Section: New Results.

83. Florian von Schilcher. The role of auditory stimuli in the courtship of Drosophila melanogaster. Animal Behaviour, 24(1):18–26, February 1976. ISSN 0003-3472. doi: 10.1016/S0003-3472(76)80095-4.

84. Daniel F. Eberl, Geoffrey M. Duyk, and Norbert Perrimon. A genetic screen for mutations that disrupt an auditory response in Drosophilamelanogaster. Proceedings of the National Academy of Sciences, 94(26):14837–14842, December 1997. doi: 10.1073/pnas.94.26.14837. Publisher: Proceedings of the National Academy of Sciences.

85. Chuan Zhou, Romain Franconville, Alexander G Vaughan, Carmen C Robinett, Vivek Jayaraman, and Bruce S Baker. Central neural circuitry mediating courtship song perception in male Drosophila. eLife, 4:e08477, September 2015. ISSN 2050-084X. doi: 10.7554/eLife.08477. Publisher: eLife Sciences Publications, Ltd.

86. Christa A. Baker, Claire McKellar, Rich Pang, Aljoscha Nern, Sven Dorkenwald, Diego A. Pacheco, Nils Eckstein, Jan Funke, Barry J. Dickson, and Mala Murthy. Neural network organization for courtship-song feature detection in Drosophila. Current Biology, 32(15): 3317–3333.e7, August 2022. ISSN 0960-9822. doi: 10.1016/j.cub.2022.06.019.

87. Steven P. Nilsen, Yick-Bun Chan, Robert Huber, and Edward A. Kravitz. Gender-selective patterns of aggressive behavior in Drosophila melanogaster. Proceedings of the National Academy of Sciences of the United States of America, 101(33):12342–12347, August 2004. ISSN 0027-8424. doi: 10.1073/pnas.0404693101.

88. Hirofumi Toda, Xiaoliang Zhao, and Barry J. Dickson. The Drosophila female aphrodisiac pheromone activates ppk23(+) sensory neurons to elicit male courtship behavior. Cell Reports, 1(6):599–607, June 2012. ISSN 2211-1247. doi: 10.1016/j.celrep.2012.05.007.

89. Judith Hoeller, Arthur Zhao, Aljoscha Nern, Edward Rogers, Sandro Romani, and Michael Reiser. The organization of visual pathways in the Drosophila brain. in prep.

90. Yoshinori Aso, Daisuke Hattori, Yang Yu, Rebecca M. Johnston, Nirmala A. Iyer, Teri-T. B. Ngo, Heather Dionne, L. F. Abbott, Richard Axel, Hiromu Tanimoto, and Gerald M. Rubin. The neuronal architecture of the mushroom body provides a logic for associative learning. eLife, 3:e04577, December 2014. ISSN 2050-084X. doi: 10.7554/eLife.04577.

91. Gerald M Rubin and Yoshinori Aso. New genetic tools for mushroom body output neurons in Drosophila. eLife, 12:RP90523, January 2024. ISSN 2050-084X. doi: 10.7554/eLife.90523. Publisher: eLife Sciences Publications, Ltd.

92. John C. Tuthill, Aljoscha Nern, Stephen L. Holtz, Gerald M. Rubin, and Michael B. Reiser. Contributions of the 12 Neuron Classes in the Fly Lamina to Motion Vision. Neuron, 79(1): 128–140, July 2013. ISSN 0896-6273. doi: 10.1016/j.neuron.2013.05.024.

93. Tanya Wolff, Mark Eddison, Nan Chen, Aljoscha Nern, Preeti Sundaramurthi, Divya Sitaraman, and Gerald M Rubin. Cell type-specific driver lines targeting the Drosophila central complex and their use to investigate neuropeptide expression and sleep regulation. eLife, 14:RP104764. ISSN 2050-084X. doi: 10.7554/eLife.104764.

94. Arnim Jenett, Gerald M. Rubin, Teri-T. B. Ngo, David Shepherd, Christine Murphy, Heather Dionne, Barret D. Pfeiffer, Amanda Cavallaro, Donald Hall, Jennifer Jeter, Nirmala Iyer, Dona Fetter, Joanna H. Hausenfluck, Hanchuan Peng, Eric T. Trautman, Robert R. Svirskas, Eugene W. Myers, Zbigniew R. Iwinski, Yoshinori Aso, Gina M. DePasquale, Adrianne Enos, Phuson Hulamm, Shing Chun Benny Lam, Hsing-Hsi Li, Todd R. Laverty, Fuhui Long, Lei Qu, Sean D. Murphy, Konrad Rokicki, Todd Safford, Kshiti Shaw, Julie H. Simpson, Allison Sowell, Susana Tae, Yang Yu, and Christopher T. Zugates. A GAL4-driver line resource for Drosophila neurobiology. Cell Reports, 2(4):991–1001, October 2012. ISSN 2211-1247. doi: 10.1016/j.celrep.2012.09.011.

95. Haojiang Luan, Nathan C. Peabody, Charles R. Vinson, and Benjamin H. White. Refined spatial manipulation of neuronal function by combinatorial restriction of transgene expression. Neuron, 52(3):425–436, November 2006. ISSN 0896-6273. doi: 10.1016/j.neuron.2006.08.028.

96. Barret D. Pfeiffer, Teri-T. B. Ngo, Karen L. Hibbard, Christine Murphy, Arnim Jenett, James W. Truman, and Gerald M. Rubin. Refinement of tools for targeted gene expression in Drosophila. Genetics, 186(2):735–755, October 2010. ISSN 1943-2631. doi: 10.1534/genetics.110.119917.

97. Kosei Sato and Daisuke Yamamoto. Mutually exclusive expression of sex-specific and non-sex-specific fruitless gene products in the Drosophila central nervous system. Gene expression patterns: GEP, 43:119232, March 2022. ISSN 1872-7298. doi: 10.1016/j.gep.2022.119232.

98. Kari Close, Yisheng He, Jennifer Jeter, Gudrun Ihrke, and Mark Eddison. Multiplex Detection of Gene Expression in the Intact Drosophila Brain Using Expansion-Assisted Iterative Fluorescence In Situ Hybridization. Journal of Visualized Experiments: JoVE, (219), May 2025. ISSN 1940-087X. doi: 10.3791/67656.

99. Alice A. Robie, Adam L. Taylor, Catherine E. Schretter, Mayank Kabra, and Kristin Branson. The Fly Disco: Hardware and software for optogenetics and fine-grained fly behavior analysis, November 2024. Pages: 2024.11.04.621948 Section: New Results.

100. Eyrun Eyjolfsdottir, Steve Branson, Xavier P. Burgos-Artizzu, Eric D. Hoopfer, Jonathan Schor, David J. Anderson, and Pietro Perona. Detecting Social Actions of Fruit Flies. In David Fleet, Tomas Pajdla, Bernt Schiele, and Tinne Tuytelaars, editors, Computer Vision – ECCV 2014, pages 772–787, Cham, 2014. Springer International Publishing. ISBN 978-3-319-10605-2. doi: 10.1007/978-3-319-10605-2_50.

101. Mayank Kabra, Alice A. Robie, Marta Rivera-Alba, Steven Branson, and Kristin Branson. JAABA: interactive machine learning for automatic annotation of animal behavior. Nature Methods, 10(1):64–67, January 2013. ISSN 1548-7105. doi: 10.1038/nmeth.2281.

